# Behavioral role of PACAP signaling reflects its selective distribution in glutamatergic and GABAergic neuronal subpopulations

**DOI:** 10.1101/2020.07.31.231795

**Authors:** Limei Zhang, Vito S. Hernández, Charles R. Gerfen, Sunny Z. Jiang, Lilian Zavala, Rafael A. Barrio, Lee E. Eiden

**Affiliations:** Department of Physiology, Faculty of Medicine, National Autonomous University of Mexico (UNAM), Mexico; Department of Complex Systems, Institute of Physics, National Autonomous University of Mexico (UNAM), Mexico; Section on Molecular Neuroscience, Intramural Research Program, NIH, Bethesda, USA; Laboratory of Systems Neuroscience, National Institute of Mental Health, Intramural Research Program, NIH, Bethesda, USA

## Abstract

The neuropeptide PACAP, acting as a co-transmitter, increases neuronal excitability, which may enhance anxiety and arousal associated with threat conveyed by multiple sensory modalities. The distribution of neurons expressing PACAP and its receptor, PAC1, throughout the mouse nervous system was determined, in register with expression of glutamatergic and GABAergic neuronal markers, to develop a coherent chemoanatomical picture of PACAP’s role in brain motor responses to sensory input. A circuit role for PACAP was tested by observing fos activation of brain neurons after olfactory threat cue in wild type and PACAP knockout mice. Neuronal activation, and behavioral response, were blunted in PACAP knock-out mice, accompanied by sharply down-regulated vesicular transporter expression in both GABAergic and glutamatergic neurons expressing PACAP and its receptor. This report signals a new perspective on the role of neuropeptide signaling in supporting excitatory and inhibitory neurotransmission in the nervous system within functionally coherent polysynaptic circuits.

## Introduction

Pituitary adenylate cyclase activating-peptide (PACAP) was first isolated and characterized from ovine hypothalamic tissue, and characterized as a peptide which stimulates cyclic AMP elevation in rat anterior pituitary cells in culture (Miyata, Arimura et al. 1989). PACAP binding to PAC1 receptors initiates signaling through multiple intracellular pathways. PACAP/PAC1 signaling is generally considered to engage Gαs, activating adenylate cyclase, and Gαq, activating phospholipase C, and leads to multiple cellular responses including increased neuronal excitability (Kawasaki, Springett et al. 1998, Emery, Eiden et al. 2013, Jiang, Xu et al. 2017, Johnson, May et al. 2019). The PACAP/PAC1 signaling pathway has consistently been related to psychogenic stress responding, and potentiation of this pathway has been linked to psychopathologies including anxiety and PTSD in human (Ressler, Mercer et al. 2011, Wang, Cao et al. 2013, Mustafa, Jiang et al. 2015). PACAP gene knock-out in the mouse results in decreased hypothalamo-pituitary-adrenal (HPA) axis activation after physical or psychogenic stress (Stroth and Eiden 2010, Tsukiyama, Saida et al. 2011), and a hypoarousal behavioral phenotype in response to psychogenic stress (Lehmann, Mustafa et al. 2013, Mustafa, Jiang et al. 2015). However, interactions within and among populations of PACAP and PAC1-expressing neurons in brain circuits mediating behavioral responses to environmental stimulation remain to be understood. This is a critical step in integrative understanding of the functional significance of PACAP-PAC1 neurotransmission.

Exploration of PACAP-containing circuits in rodent central nervous system (CNS) has been based on reports of the distribution of PACAP peptide and mRNA, and on expression from reporter genes under the control of a PACAP promoter transgene (Hannibal 2002, Condro, Matynia et al. 2016, Koves 2016) or knocked-in to the PACAP gene itself (Krashes, Shah et al. 2014). Hannibal reported the anatomical distribution of PACAP projection fields and cell groups in rat CNS employing immunohistochemistry (IHC) and *in situ* hybridization (ISH), using radio-labeled riboprobes, in a rigorous study. However, due to the paucity of PACAP in cell bodies, dendrites and axons compared to nerve terminals, peptide IHC has not provided more definitive PACAP chemoanatomical circuit identification in rodent brain. Similarly, ISH with radiolabeled riboprobes, while identifying PACAP-positive cell bodies, lacks the resolution to identify the co-transmitter phenotypes and precise microanatomical features of these cell groups. Thus, heterogeneity of PACAP-containing neurons within and between brain regions, both with respect to cell type and accurate regional boundaries could not be discerned. More recently, lesion as well as micro-infusion approaches have tentatively identified some PACAP containing projection systems (Miles, Thrailkill et al. 2017). Nevertheless, an essential function for PACAP in the context of neurotransmitter action within one or more brain behavioral circuits, and consistent with the cellular and post-synaptic actions of PACAP, has not yet emerged. A systematic analysis with accuracy at the level of cellular co-phenotypes, and with anatomical resolution to the level of sub-nuclei within CNS, is essential to complete this task.

## Results

We have studied PACAP and PAC1 mRNA co-expression with VGLUT1, VGLUT2 and VGAT mRNAs in the mouse brain, with precise region and subfield identification, using a sensitive dual ISH (DISH) method. Figures 1 (PACAP mRNA) and 2 (PAC1 mRNA) show examples using this method, which unambiguously labels the co-expression of two mRNAs at the single cell level for light microscopic examination. At the light microscopic level, facile low- and high-magnification switching allows detailed serial high-power images to be located in a global histological context for precise delineation of anatomical regions/subfields as well as their rapid photo-documentation. Single cell co-expression of two mRNA targets can be clearly observed by light microscopy with both low and high magnification (anatomical details for each panel, *vide infra*).

**Figure 1.**
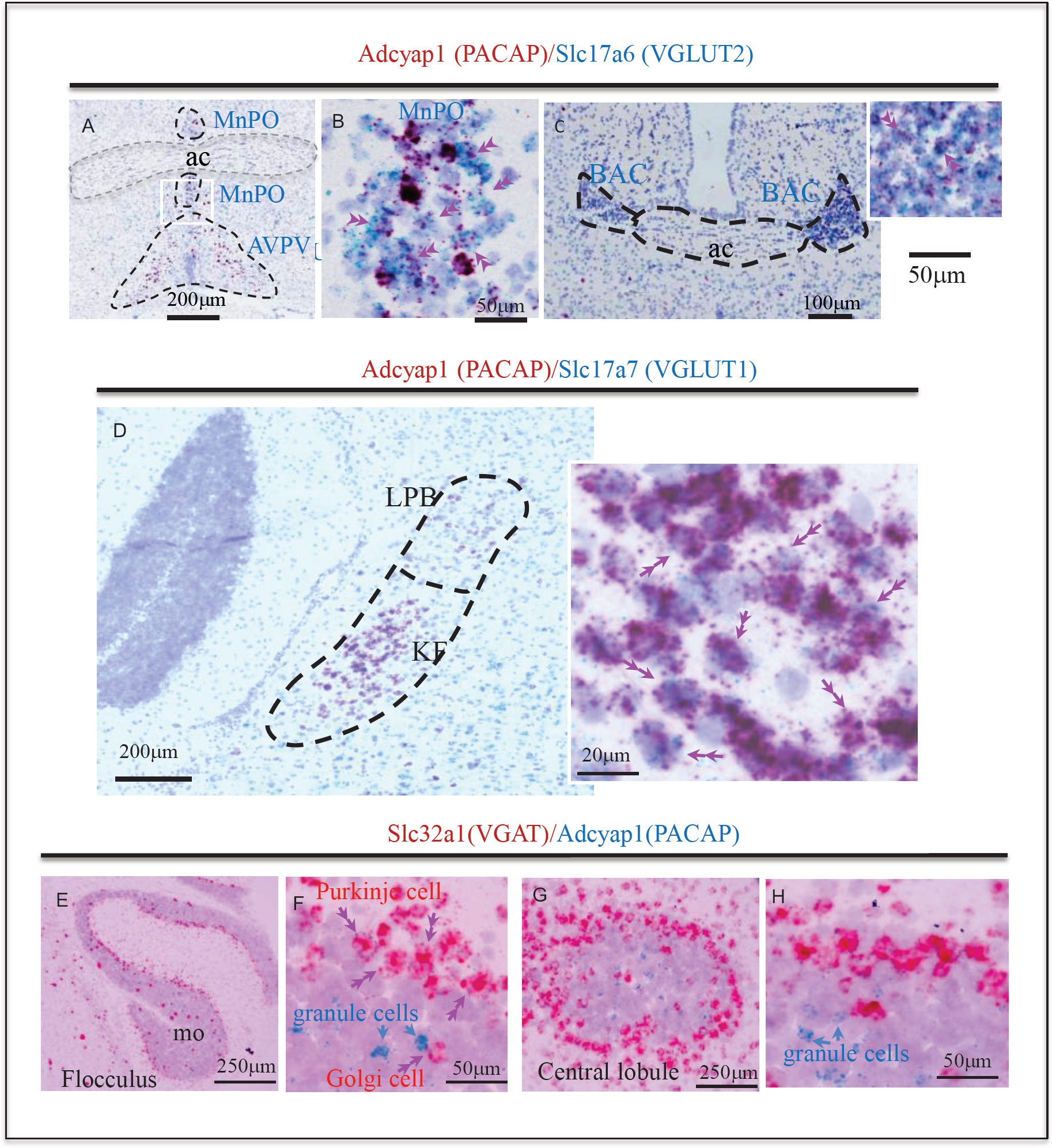
Examples of histological samples using the sensitive dual ISH method that can label unambiguously the co-expression of two RNAs at single cell level for light microscope examination. A-C: Adcyap1 (RNA encoding for PACAP) co-expression with Slc17a6 (RNA encoding for VGLUT2) in medial preoptic nucleus (MnPO**) (A, B)** and bed nucleus of anterior commissure (BAC) (C). Single cell co-expression feature can be clearly observed in the high magnification photomicrographs (B and inset of C). This feature is better appreciated on the cells in which Adcyap1 is weakly expressed (week staining), so the Slc17a6 staining can be clearly seen as independent dots (indicated by double arrows). D: brain stem Koelliker Fuse (KF) nucleus of the parabrachial complex is another main PACAP-expressing nucleus described in the literature (for instance (Hannibal 2002)). The vesicular transporter RNA strongly co-expressed with Adcyap1 here is Slc17a7 (RNA encoding VGLUT1). Figure E-H show two cerebellar regions, paraflocculus (E and F) and central (G and H), low and high magnification respectively, where the Adcyap1 expression is higher than rest of regions. Purkinje cells are the main GABAergic (expressing Slc32a1, RNA encoding VGAT) PACAP containing, distributed in all regions of cerebellar cortex. Some Golgi cells in granule cell layer of paraflocculus and central regions also co-expressed Adcyap1 and Slc32a1 (indicated with double pink arrowheads. In these two cerebellar regions, some granule cells also expressed Adcyap1 (indicated with single blue arrows).

### 1. Comprehensive DISH mapping of PACAP co-expression with VGLUT1, VGLUT2 and VGAT throughout mouse brain reveals an extensive distribution and diversity of cell types

Table 1 describes the distribution, cell types and relative expression strength, within 163 identified PACAP mRNA-positive cell groups/subfields co-expressing vesicular transporters, organized hierarchically by grouping the regions according to their embryonic origins. These include 53 regions derived from cortical plate, 6 regions derived from cortical subplate, 11 regions within cerebral nuclei, 19 regions in thalamus, two regions in epithalamus, 23 regions in hypothalamus, 17 regions in midbrain, 14 regions in pons, 18 regions in medulla and five regions in cerebellum. To compare with the previous comprehensive report for PACAP distribution in rat brain published in 2002 (Hannibal 2002), a column containing the data published previously in rat is displayed in blue. Most of the regions described as PACAP-expressing in the rat were also found positive in our study in mouse, albeit strength of expression in several regions differs substantially between the two rodent species. Eleven regions that were reported negative, labeled as ‘-’ from original publication, were found positive with this sensitive DISH method (indicated in the table). An additional 108 regions, which were not reported in detail in the previous paper (labeled in the table as “n/r”), were found to co-express PACAP and either a glutamate or GABA vesicular transporter.

**Table 1.**
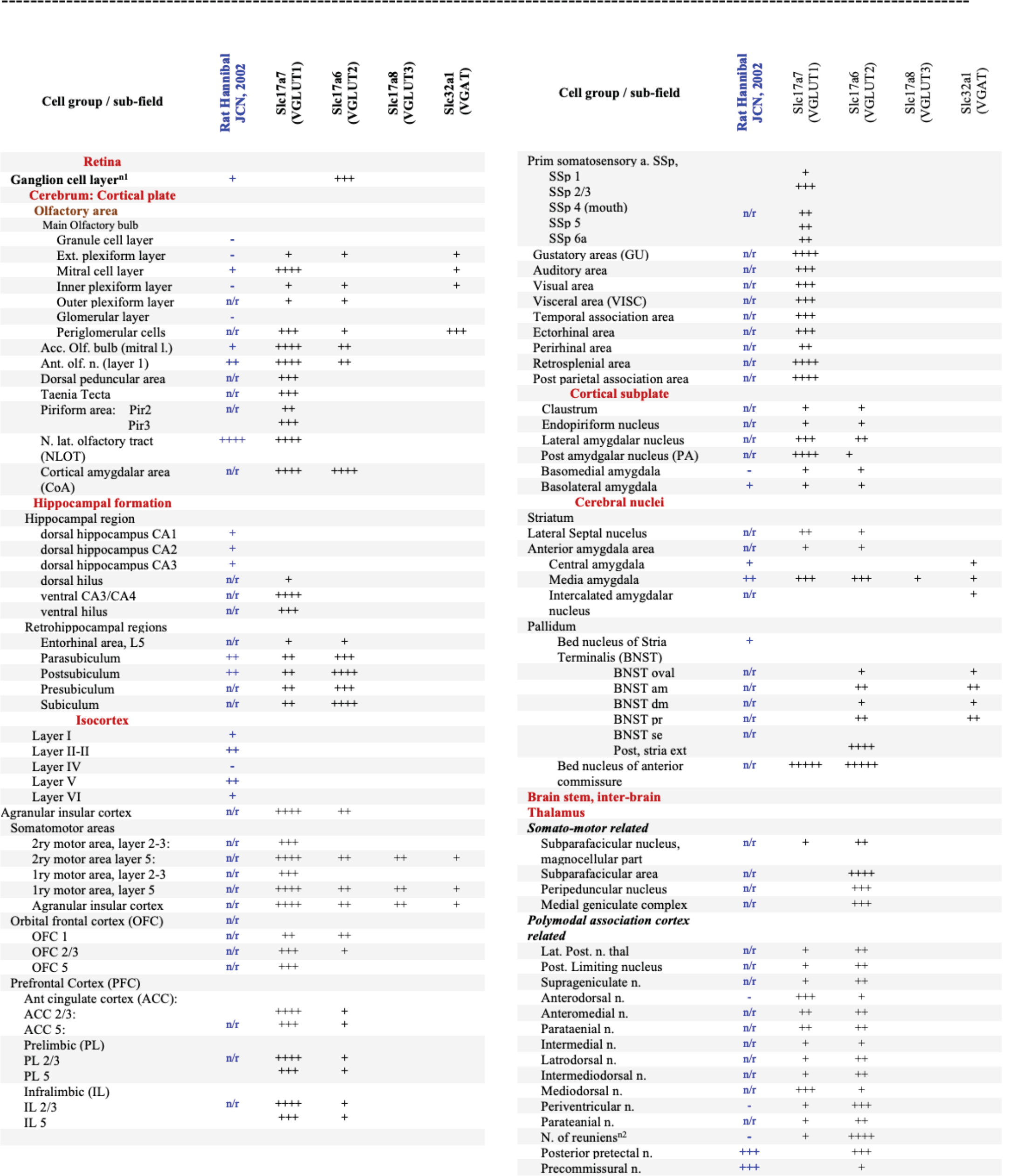
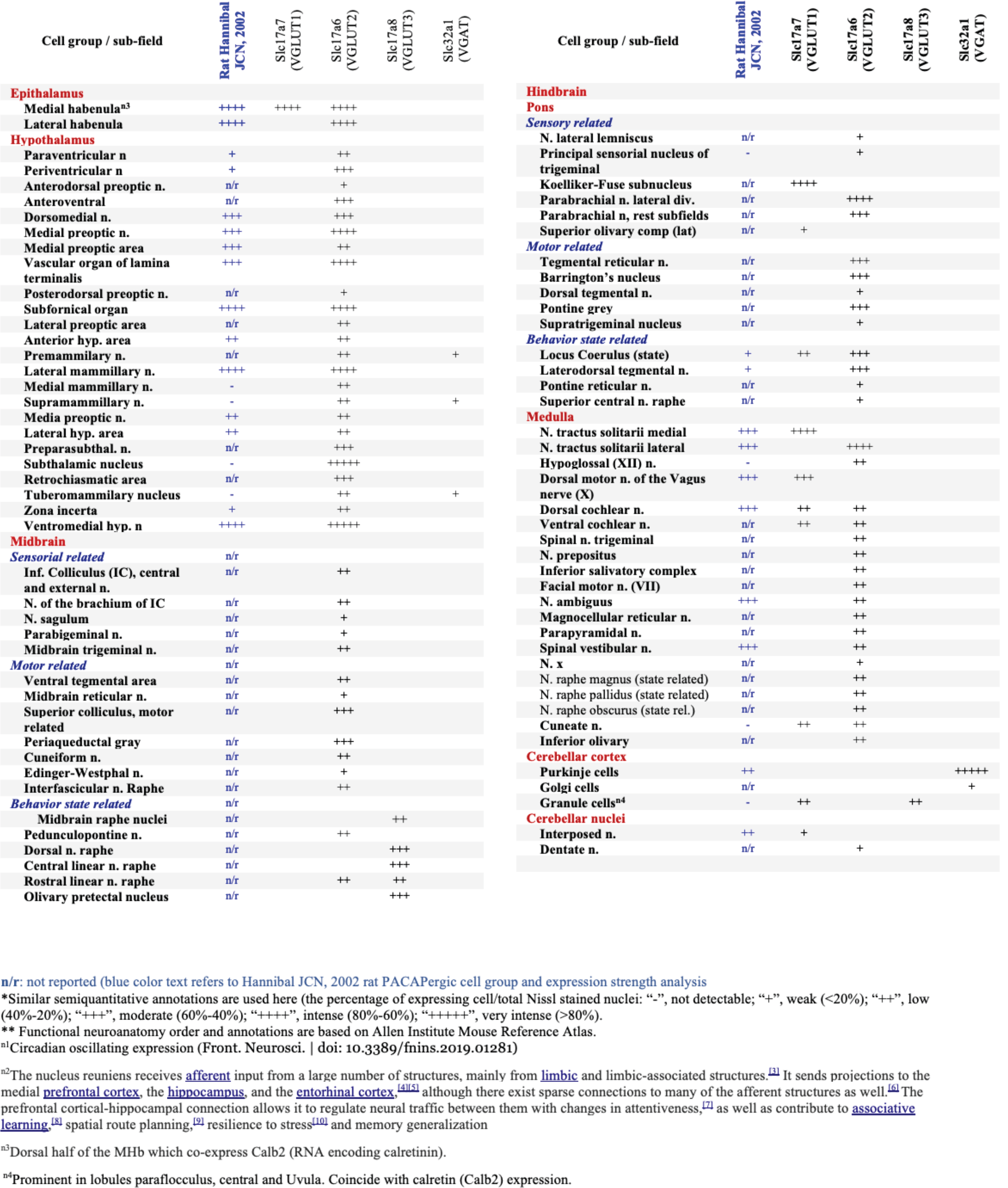
Distribution, cell types and strength of main **PACAPergic cell groups** in mouse brain with comparison to rat brain reported by Hannibal, JCN, 2002*

A whole brain mapping of PACAP mRNA expression with relevant brain regions/subfields co-expression features is presented in the supplemental information (SI) Fig. 1.

### 2. Mapping of PAC1 mRNA co-expression with mRNAs for PACAP, VGLUT1, VGLUT2, VGAT suggests the PACAP-PAC1 system can function in *autocrine* and *paracrine* modes

PAC1 mRNA-expression was studied in 150 mouse brain regions. In figure 3, panels A-F, we show the semi-quantitative expression levels of PAC1 mRNA, based on microscopic observation as different intensities of blue shading. PACAP mRNA expression was also symbolized with either red or green dots (VGAT vs. VGLUT co-expression) in corresponding regions. Contrasting with the discrete expression of PACAP mRNA, the PAC1 mRNA expression was diffuse, and widespread. PAC1 cells co-expressed Slc17a7 (VGLUT1 mRNA) in the temporal hippocampus (Fig. 2 panel A and B), anterior cingulate area (ACA, panel C) and bed nucleus of anterior commissure (BAC) and Scl32a1 (VGAT mRNA) in pallidum and striatum structures (Fig. 2, D-G and Tab. 2). Almost all the PACAP mRNA-expressing neurons we studied co-expressed PAC1 mRNA (Fig. 3, G, from ACA and H from medial preoptic nucleus, MnPO). Besides, most neurons neighboring PACAP positive cells also expressed PAC1 receptors (single arrows). These observations suggest that the PACAP/PAC1 pathway uses *autocrine* and *paracrine* mechanisms in addition to classical neurotransmission through axon innervation and transmitter co-release.

**Figure 2.**
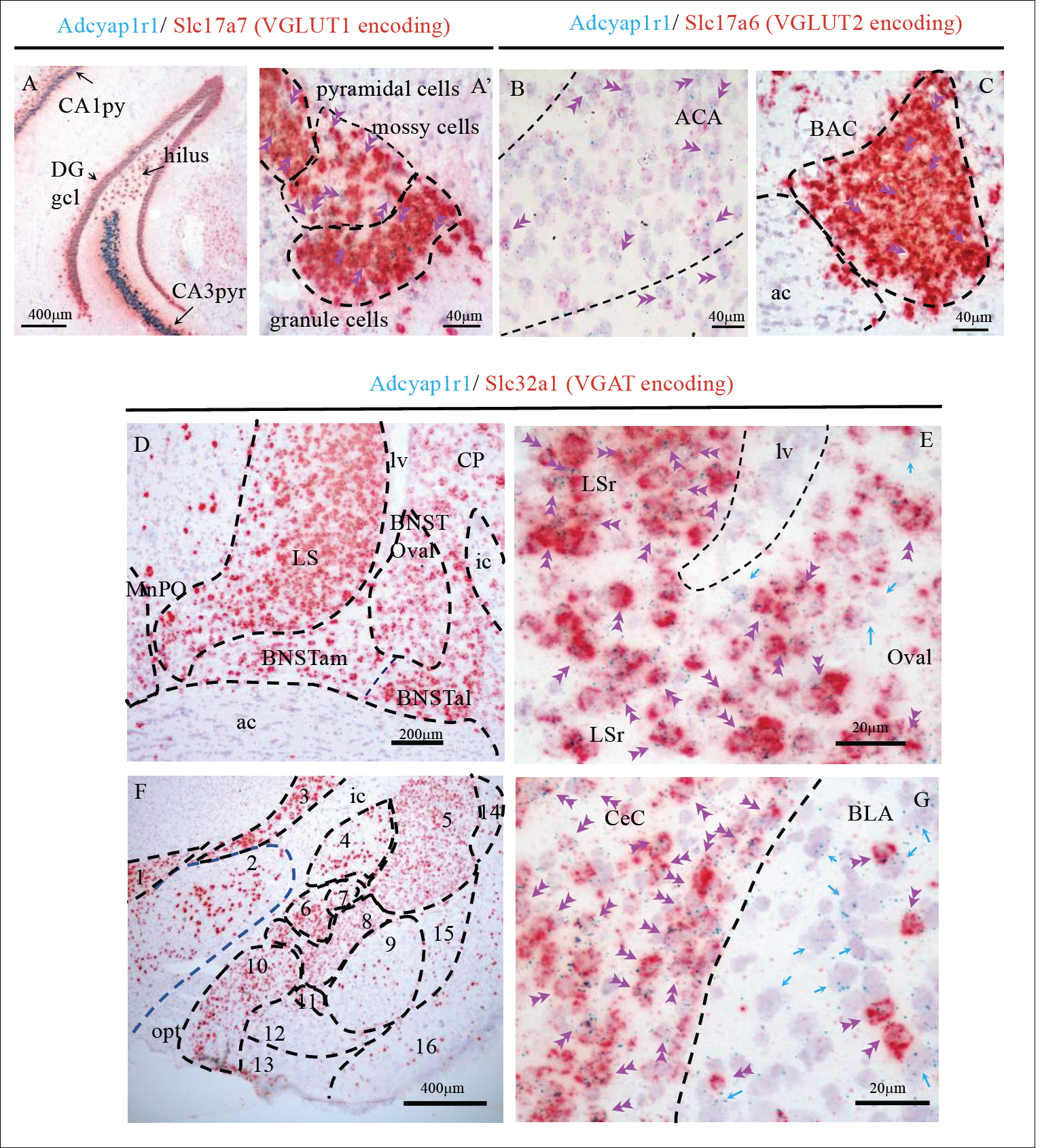
Examples illustrating Adcyap1r1-co-expression with glutamatergic (Slc17a7-VGLUT1 and Slc17a6-VGLUT2 expressing) and GABAergic (Slc32a1 expressing) neurons in cortical and subcortical regions. A and A’: temporal hippocampal formation where the Adcyap1r1 was strongly expressed in the principal neurons (pyr: pyramidal layer and DGgcl: dentate gyrus granule cell layer) as well as the VGLUT1+ mossy cells in the hilar region. B. Adcyap1r1-co-expression with Slc17a6 (VGLUT2 encoding) in anterior cingulate area (ACA) and D in the bed nucleus of anterior commissure (BAC). Regarding the GABAergic neurons expressing Adcyap1r1, the structures in the striatum and pallidum host the strongest expressing structures. Panel D shows the lateral septum (LS), bed nucleus of stria terminalis (BNST) in its three divisions, anteromedial (BNSTam), antero-lateral (BNSTal) and oval (BNSToval), as well as caudate-putamen (CP) with highest Adcyap1r1 expression. Panel E shows high magnification photomicrograph where green dots, Adcyap1r1 labeling, are mostly overlapped with red staining (Slc32a1, VGAT expression), double pink arrowheads, while some single-expressed cells are also observed (single green arrows). Panel F shows amygdaloid complex and neighboring regions where Adcyap1r1 was strongly expressed in the GABAergic cell populations; G: high magnification photomicrograph showing that the Adcyap1r1 is exclusively expressed in Slc32a1 (VGAT) expressing neurons in the CeC, while in the BLA it was expressed in the sparsely distributed GABAergic neurons as in most of the non-VGAT expressing neurons. 1. zona incerta of hypothalamus; 2. lateral hypothalamic area; 3: reticular nucleus of the thalamus; 4. globus pallidus; 5. caudate-putamen; 6: central amygdalar nucleus, medial part (CeM); 7: lateral part (CeL); 8: capsular part (CeC); 9: basolateral amygdalar nucleus (BLA) 10: medial amigdalar nucleus; 11: intercalated nucleus of the amygdala; 12: basomedial nucleus of the amygdala; 13: cortical amygdalar area; 14: dorsal endopiriform; 15: ventral endopiriform; 16: piriform area. Fiber tracts: sm: stria medullaris; ac: anterior commissure; ic: internal capsule; opt: optic tract; lv: lateral ventricle; ic: internal capsule.

**Figure 3.**
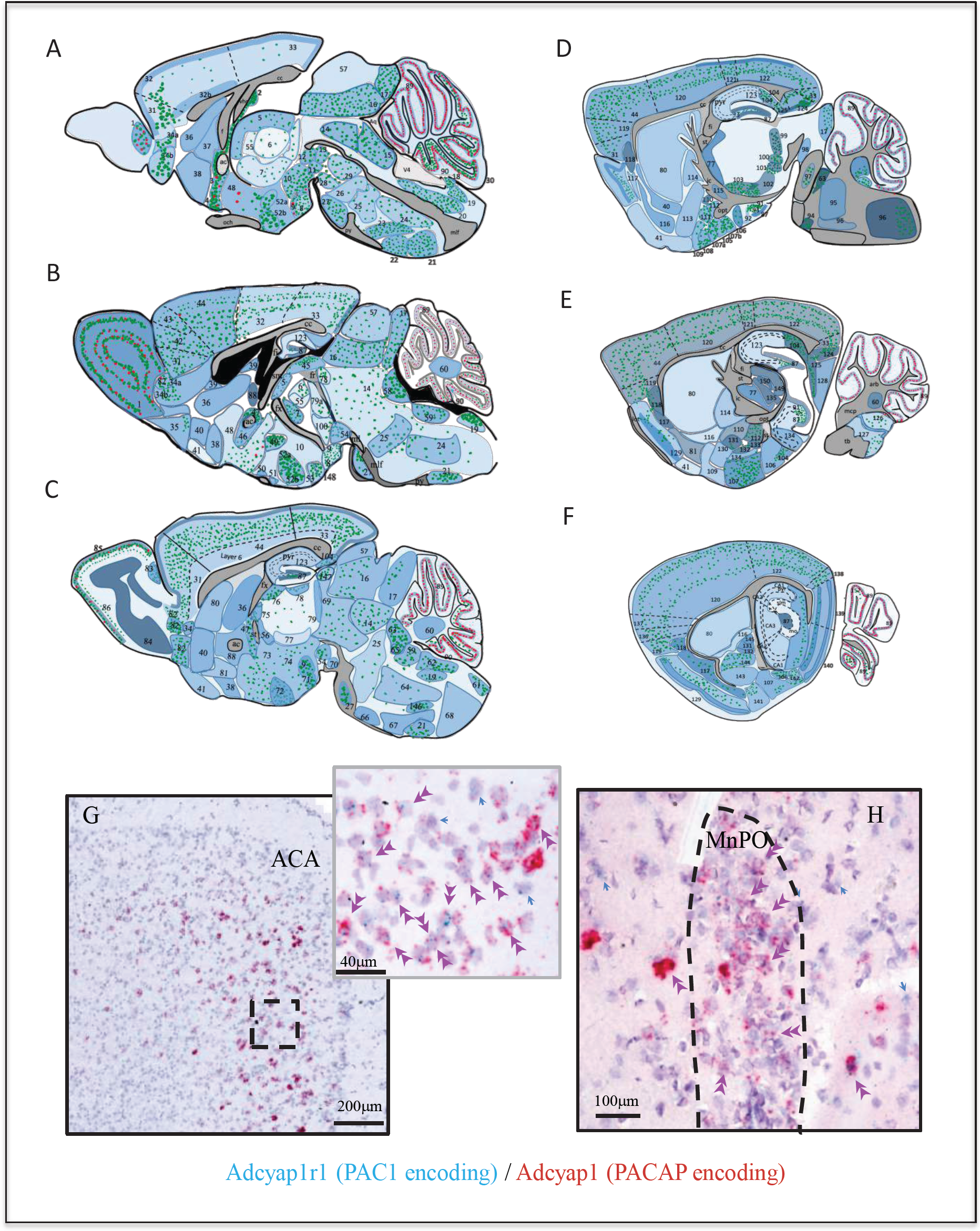
Adcyap1r1 expression assessment in relation to Adcyap1 expression suggests the PACAP-PAC1 system uses autocrine, paracrine and neuroendocrine modes for signal transduction. A-F: Mapping of Adcyap1r1 (RNA encoding Pac1) expression (symbolized by the intensity of blue shading) in six septo-temporal planes in relation to main PACAPergic brain nuclei and subfields, based on microscopic observations. Green and red dots represent Adcyap1 expressing neurons of glutamatergic (vesicular glutamate transporter expressing) or GABAergic (VGAT expression) nature, respectively. Shaded regions with different blue intensity symbolize the strength of Adcyap1r1. *Figure 3 continued on next page* 1. main olfactory bulb; 2. subfornical organ; 3. median preoptic n.; 4. vascular organ of lamina terminalis; 5. paraventricular n. of the thalamus; 6. mediodorsal n. of the thalamus; 7. n. reuniens; 8. mammillary n.; 9. supramammilary n.; 10. posterior hypothalamic n.; 11. interfascicular n. raphé; 12. Edinger-Westphal n.; 13. rostral linear n. raphé; 14. periaqueductal grey; 15. dorsal raphé n.; 16. superior colliculus, motor related; 17. inferior colliculus; 18. area postrema; 19. n. tractus solitarius; 20. XII hypoglossal n.; 21. inferior olivary complex; 22. n. raphé pallidus; 23. n. raphé magnus; 24. gigantocellular reticular n.; 25. pontine reticular n; 26. trigeminal reticular n.; 27. pontine grey; 28. interpeduncular n.; 29. central linear n. raphé; 30. uvula of cerebellum; 31. orbital area; 32. anterior cingulate area, dorsal, layer 1; 32b. anterior cingulate area, dorsal, layer 6; 33. retrosplenial cortex; 34a. dorsal peduncular area; 34b. taenia tecta; 35. taenia tecta, ventral; 36. lateral septal n., rostral; 37. medial septal n.; 38. diagonal band nucleus; 39. lateral septal n., caudal; 40. n. accumbens; 41. olfactory tubercle; 42. infralimbic area; 43. prelimbic area; 44. secondary motor area; 45. medial habneula; 46. bed nucleus of stria terminalis, oval n.; 47. bed nucleus of anterior commissure; 48. medial preoptic area; 49. hypothalamic paraventricular n.; 50. suprachiasmatic nucleus; 51. supraoptic nucleus; 52a. dorsomedial hypothalamic n; 52b. ventromedial hypothalamic n.; 53. periventricular hypothalamic n., posterior part; 54. ventral tegmental area; 55. anteromedial n.; 56. parataenial n.; 57. superior colliculus, sensory related; 58. pontine central grey; 59. medial vestibular n.; 60. cerebellar nuclei: fastigial n.; 61. cuneate n.; 62. spinal vestibular n.; 63. parabrachial n.; 64. intermediate reticular n.; 65. locus coeruleus; 66. superior olivary complex; 67. magnocellular reticular n.; 68. medullary reticular n.; 69. midbrain reticular n.; 70. substantia nigra; 71. lateral mammillary n.; 72. ventrolateral hypothalamic n.; 73. lateral preoptic area; 74. posterior hypothalamic area; 75. anteroventral n. of thalamus; 76. anterodorsal n. of the thalamus; 77. reticular n. of the thalamus; 78. precommisural n.; 79. parafascicular n.; 80. caudoputamen; 81. substantia innominata; 82. anterior olfactory n.; 83. accessory olfactory bulb; 84. main olfactory bulb, granule layer; 85. main Olfactory bulb, glomerular layer; 86: main olfactory bulb, mitral layer; 87. hippocampal formation, granule cell layer of dentate gyrus; 88. bed n. of stria terminalis; 89. Purkinje cell layer of cerebellum; 90. Granular layer of cerebellum; 91. hippocampal formation, ventral CA3c; 92. ventral CA3; 93. ventral hilus; 94. motor n. of trigeminal nerve; 95. principal sensory n. of trigeminal; 96. spinal nucleus of trigeminal; 97. n. of lateral lemniscus; 98. parabigeminal nucleus; 99. medial geniculate complex; 100. subparafascicular nucleus; 101. peripeduncular nucleus; 102: zona incerta; 103. subthalamic nucleus; 104. subiculum; 105. medial amydgala, posteroventral part; 106. posterior amygdala nucleus; 107a. cortical amygdala anterior part; 107b. cortical amygdala posterior part; 108. n. of the lateral olfactory tract; 109. anterior amygdala area; 110. Central amygdala medial part; 111. medial amydgala, anterodorsal part; 112. medial amygdala, posterodorsal part; 113. Substantia innominata; 114. globus pallidus, external part; 115. globus pallidus, internal part (entopeduncular nucleus); 116. fundus of striatum; 117. endopiriform n.; 118. claustrum; 119. Agranular insular cortex; 120. somatosensory area; 121. posterior parietal association area; 122. visual area; 123. dorsal hippocampus proper; 124. postsubiculum; 125. presubiculum; 126. dorsal cochlear nucleus; 127. ventral cochlear nucleus; 128. parasubiculum; 129. piriform area; 130. anterior amygdala area; 131. central amygdala, capsular part; 132. intercalated amygdalar nucleus; 133. basomedial amygdala nucleus; 134. ventral hippocampus, CA2; 135. lateral geniculate complex, ventral parte; 136. gustatory area; 137. visceral area; 138. ectorhinal area; 139. entorhinal area, lateral part; 140. entorhinal area, medial part; 141. piriform-amygdalar area; 142. postpiriform transition area; 143. basolateral amygdala; 144. lateral amygdala. 145. Central amygdala, lateral part. 146. n. ambiguus; 147. Olivary pretectal nucleus; 148: premammilary n.; 149. lateral geniculate complex, dorsal part; 150. intergeniculate leaflet; 151. cerebellar cortex, flocculus, granule cell layer; 152. cerebellar cortex, ansiform lobule, granule cell layer. Aq: aqueduct; och: optic chiasm; v4: forth ventricle; mlf: medial longitudinal fasciculus; cc: corpus callosum; vhc: ventral hippocampus commissure; fi/fx: fimbria/fornix; py: pyramidal layer; lot: lateral olfactory tract, mcp: middle cerebellar penducle; st: stria terminalis; opt: optic tract; ic: internal capsule; tb: trapezoid body; arb: arbor vidae. G and H: Examples illustrating *autocrine* and *paracrine* features of PACAP-Pac1 signaling: Adcyap1r1-co-expression in cortical and subcortical regions’ PACAP containing (Adcyap1 expressing) neurons. G: anterior cingulate area (ACA) in prefrontal cortex and H: median preoptic nucleus (MnPO). Double arrowheads indicate co-expression and blue arrows indicate the Adcyap1r1 expressing neurons which are not Adcyap1 expressing but were adjacent to them.

PACAP binds to two other G protein–coupled receptors highly related to PAC1, called VPAC1 (Vipr1), and VPAC2 (Vipr2) (Harmar 2001). In all the regions where PAC1 mRNA was expressed, the expression of mRNA for either or both VIP receptors was also found along with VGLUT1, VGLUT2 or VGAT mRNAs (data not shown here). Vipr1 (VPAC1) expression was widespread, like PAC1 mRNA, in cerebral cortex, hippocampal formation (prominently in the mossy cells), structures derived from cerebral subplate and cerebral nuclei, as well as hypothalamus. The cerebellar cortex and deep cerebellar nuclei moderately expressed, and the Purkinje cells strongly expressed Vipr1; Vipr2 (VPAC2) was observed to be strongly and selectively expressed in the MOB, the mitral and granule layers, the BNSTov, CeA, lateral division, the SCN, ventral anterior, posterior, posterior medial and lateral geniculate nuclei of thalamus, cranial nerve nuclei III, V, VII, pontine reticular nucleus, superior olivary complex, nucleus raphe pontis, and paragigantocellular reticular nucleus.

### 3. Distribution and glutamatergic/GABAergic co-expression of PACAP/PAC1 suggests a broad function for PACAP signaling in sensorimotor processing system(s)

#### 3.1 Retina

Retinal ganglion cells (RGC) have been reported to express PACAP at various levels of abundance previously in the literature. In rat, it was reported as a low level (“+”) (Hannibal 2002) within the RGC population. CD1 mice were reported to express PACAP in around 40% of RGCs (Kawaguchi, Isojima et al. 2010) and in PACAP-EGFP mice, reporter gene expressed was reported to be low (“+”) (Condro, Matynia et al. 2016). With the DISH method employed here, we found a higher percentage of retina ganglion cells co-expressing Adcyap1 and Slc17a6 (mRNAs encoding PACAP and VGLUT2) than previously reported. Expression levels of PACAP mRNA oscillate daily from 50% to 80% with highest levels during subjective night (SI, Fig. 1-A3(Lindberg, Mitchell et al. 2019)).

#### 3.2 Cerebral cortex: structures derived from cortical plate

##### 3. 2. 1 Olfactory area

High levels of PACAP expression in the olfactory area have been previously reported (Hansel, May et al. 2001). Here we report the types of olfactory area PACAP mRNA-expressing neurons in detail. In the *main olfactory bulb* (MOB), PACAP mRNA was strongly expressed with either VGLUT1 or VGAT mRNA in the mitral cell layer and periglomerular cells (see the supplemental information SI, Fig. 1-A, A1 and A2). Other cell types in the external and internal plexiform cell layers expressed PACAP mRNA at low levels with mixed glutamate/GABA molecular signatures (see Tab. 1). In contrast, in the *accessory olfactory bulb* (AOB), PACAP mRNA was mainly expressed in the mitral cell layer with co-expression of both VGLUT1 and 2 mRNAs. Other olfactory areas strongly expressing PACAP and VGLUT1 mRNAs were AON (layer 1, L1), TT, DPA, Pir (L3), CoA (L2). The NLOT (L3) co-expressed PACAP/VGLUT2 mRNA (see Tab. 1 and SI Fig. 1 for more details).

##### 3. 2. 2 Isocortex

PACAP’s role in isocortex has in general been little studied (Zhang and Eiden 2019). Moderate expression of PACAP mRNA transcripts were initially reported in the cingulate and frontal cortices, with lower concentrations found in other neocortical areas using radiolabeled riboprobe ISH (Mikkelsen, Hannibal et al. 1994). Hannibal subsequently reported that PACAP mRNA-expressing cells were observed mainly in layers 1–3 and layers 5–6, and PACAP-IR nerve fibers in all layers of the cerebral cortex; however, no detailed information about the differential expression levels across cortical regions was presented (Hannibal 2002).

In our study, we found the strongest expression of PACAP mRNA in isocortex to be in the frontal pole of the telencephalon, including prelimbic, infralimbic and anterior cingulate area, dorsal and orbital area. Approximately 80% of the neuronal population of the layer 2 and layer 5 co-expressed PACAP and VGLUT1 mRNAs. A significant population of PACAP mRNA-expressing cells in the layer 5 co-expressed VGLUT2 mRNA or VGAT mRNA (Tab. 1 and SI Fig. 1, panels B and B1).

In the primary and secondary motor cortices (MOp and MOs, respectively), PACAP mRNA was found expressed in layer 2/3 and layer 5. This pattern was also observed in somatosensory, gustatory, auditory, visual, visceral, temporal association, ectorhinal, perirhinal, retrosplenial, and post-parietal association areas (see Tab. 1 for more cortical area expression and strength and SI Fig. 1, corresponding panels for expression patterns).

PAC1 mRNA expression in neocortex was widespread, with more than 80% of neurons in layers 2-6 expressing PAC1 mRNA (see SI-Tab. 1). Across cortical regions, distribution among cortical layers was quite similar, except that in the ACA and the entorhinal cortex layers 2-3 and layers 5-6 showed highest expression levels (Fig. 3, panels B and F). As approximately 20% of cortical neurons are GABAergic (Petilla Interneuron Nomenclature, Ascoli et al. 2008), we tested the three main GABAergic cell types in the cortical regions, finding 100% of somatostatin (Sst), parvalbumin (PV) and corticotropin releasing hormone (CRH) neurons co-expressed PAC1 (SI Fig. 3, F, G, H).

##### 3. 2. 3 Hippocampal formation

In the mouse **dorsal (septal pole) hippocampal formation**, in contrast to data obtained in rat (SI Fig. 2) and from the PACAP-EGFP transgenic reporter mouse, we did not find PACAP mRNA expressing cells in cell body layers regions, *i*.*e*. the pyramidal cell layer nor in the granule cell layer (GCL) of the dentate gyrus (DG), as previously reported (Hannibal 2002, Condro, Matynia et al. 2016). PAC1-mRNA expression was observed to be sparse in CA subfields and was selectively expressed in VGAT-mRNA expressing cells (SI Fig. 5, panel A and B). In contrast, the DG-GCL had the highest expression level of PAC1-mRNA, among all brain regions, in both VGLUT1 and VGAT mRNA expressing cells (Fig. 2, panel A and SI Fig. 3 A and C). In DG hilar region (polymorphic layer), we observed few cells co-expressed PACAP and VGLUT1 mRNA (Fig. 4-A, SI Fig 1 panel I_7_). These were *mossy cells* co-expressing calretinin mRNA (Calb2) (Fig. 4-B). The PACAP mRNA-expressing mossy cell’s quantity increased in the caudo-temporal direction. This population was also described in the previous reports (Hannibal 2002, Condro, Matynia et al. 2016), however, without identification of cell type, as reported here.

**Figure 4.**
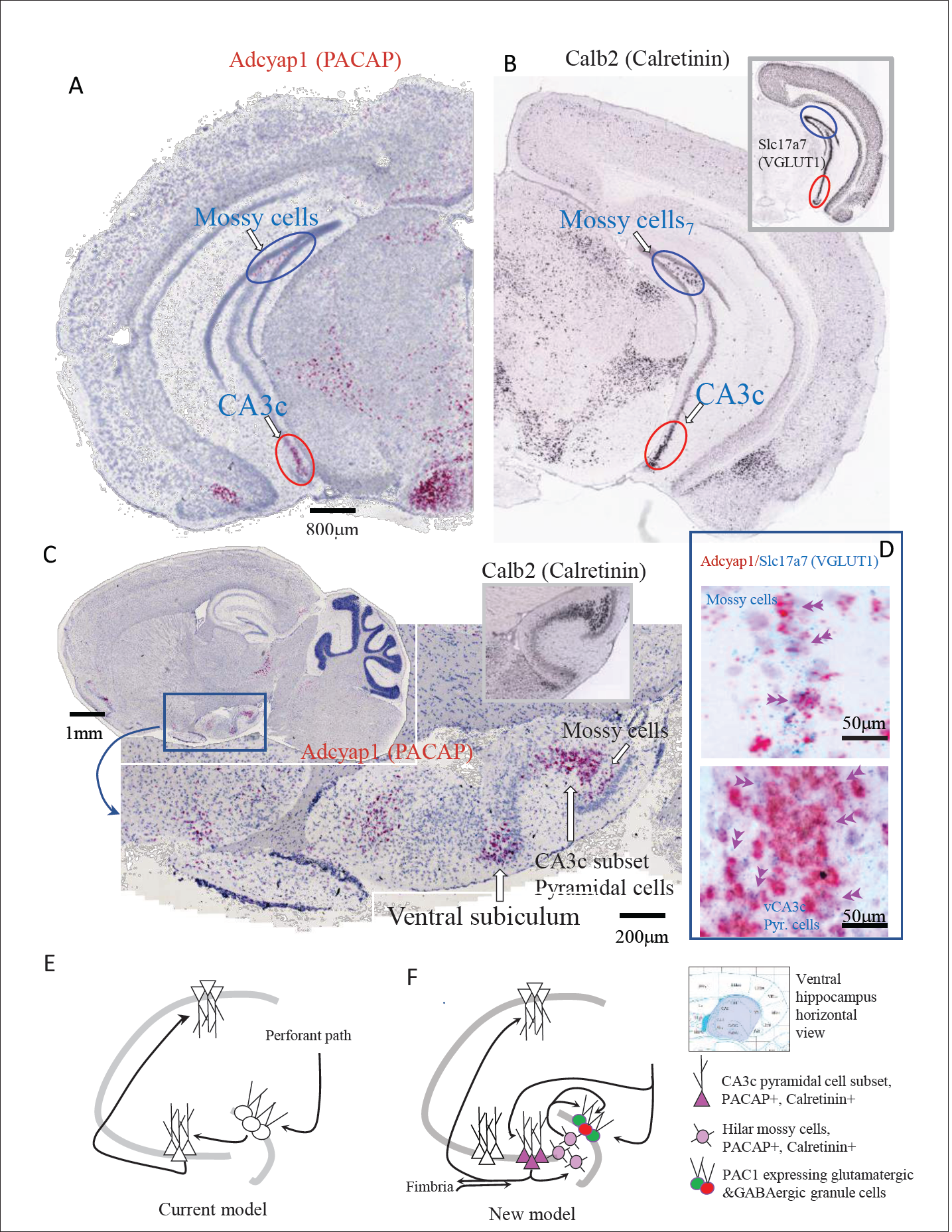
Ventral (temporal pole) hippocampus CA3c hosted a newly identified subset of pyramidal neurons distinguished by its molecular signatures of VGLUT1, PACAP and calretinin expression. Low magnification bright field coronal (A) and sagittal (C) whole-brain sections with ISH (RNAscope® 2.5 High Definition (HD) Red Assay), showing the selective expression of mRNA of PACAP (Adcyap1) in a subset of CA3c of the temporo-ventral pole of hippocampus (red ovals circumscribed regions in A and C). Hilar mossy cells in the dorsal (A) and ventral (C) hippocampus, also Adcyap1 expressing, are circumscribed by blue ovals. B and inset show the corresponding coronal sections in low magnification of Calb2 (calretinin) and slc17a7 (VGLUT1) taken from Allen Brain Atlas (Ng et al, 2009, An Anatomic Gene Expression Atlas of the Adult Mouse Brain-PMID 19219037) where CA3c subset and hilar mossy cells are indicated with red and blue ovals, respectively). C-inset corresponds calretinin mRNA expression in the same hippocampus sagittal region of C. Both the subset of CA3c pyramidal neurons and the mossy cells co-expressed VGLUT1 mRNA (Slc17a7) and Adcyap1 RNAscope® 2.5 HD Duplex Assay –(D). The “trisynaptic-centric” (E) vs “CA3-centric” (F) view of hippocampal information processing, where the newly identified CA3c subset of, PACAP & calretinin containing glutamatergic neurons are presented in dark pink triangles and the mossy cells are in light pink circles. The PAC1 expressing granule cells (green) and interneurons (red) in the granule cell layer are symbolized with pink circle. Chartings were based on ventral pole of hippocampus (shaded region of atlas segment *(Paxinos Mouse brain)*, where this chemically distinct subset of CA3c pyramidal neurons was identified. Circuits were modified from (Scharfman 2007), with adaptation to the new finding from this study.

**Figure 5.**
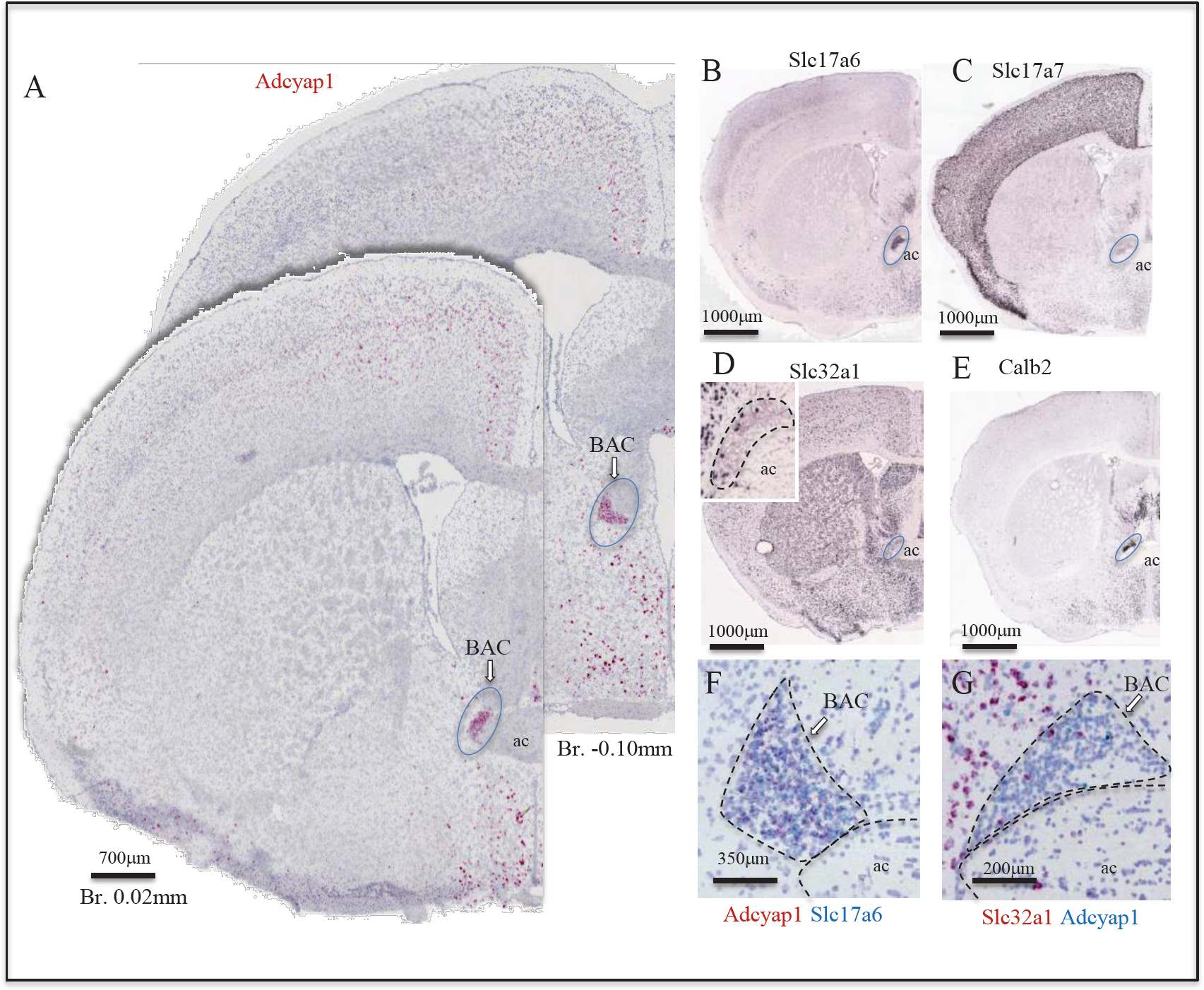
Bed nucleus of anterior commissure (BAC): a prominent glutamatergic-PACAPergic nucleus chemo-anatomically identified. A: two coronal sections at Bregma 0.02 mm and -0.10 mm of mouse brain showing Adcyap1 ISH (RNAscope® 2.5 High Definition (HD) Red Assay) expressing BAC (ac: anterior commissure). Panels B-E are low magnification photomicrographs taken from Allen Brain Atlas (Ng et al, 2009, An Anatomic Gene Expression Atlas of the Adult Mouse Brain-PMID 19219037) showing the Slc17a6 (VGLUT2, B), Slc17a7 (VGLUT1, C), Slc32a1 (VGAT, D and inset), Calb2 (calretinin, E) expressed in BAC. The Adcyap1 expressing neurons were densely packed and co-expressed Slc17a6 (F), but we also observed co-expression within the Slc32a1 expressing cells (G).

**Figure 6.**
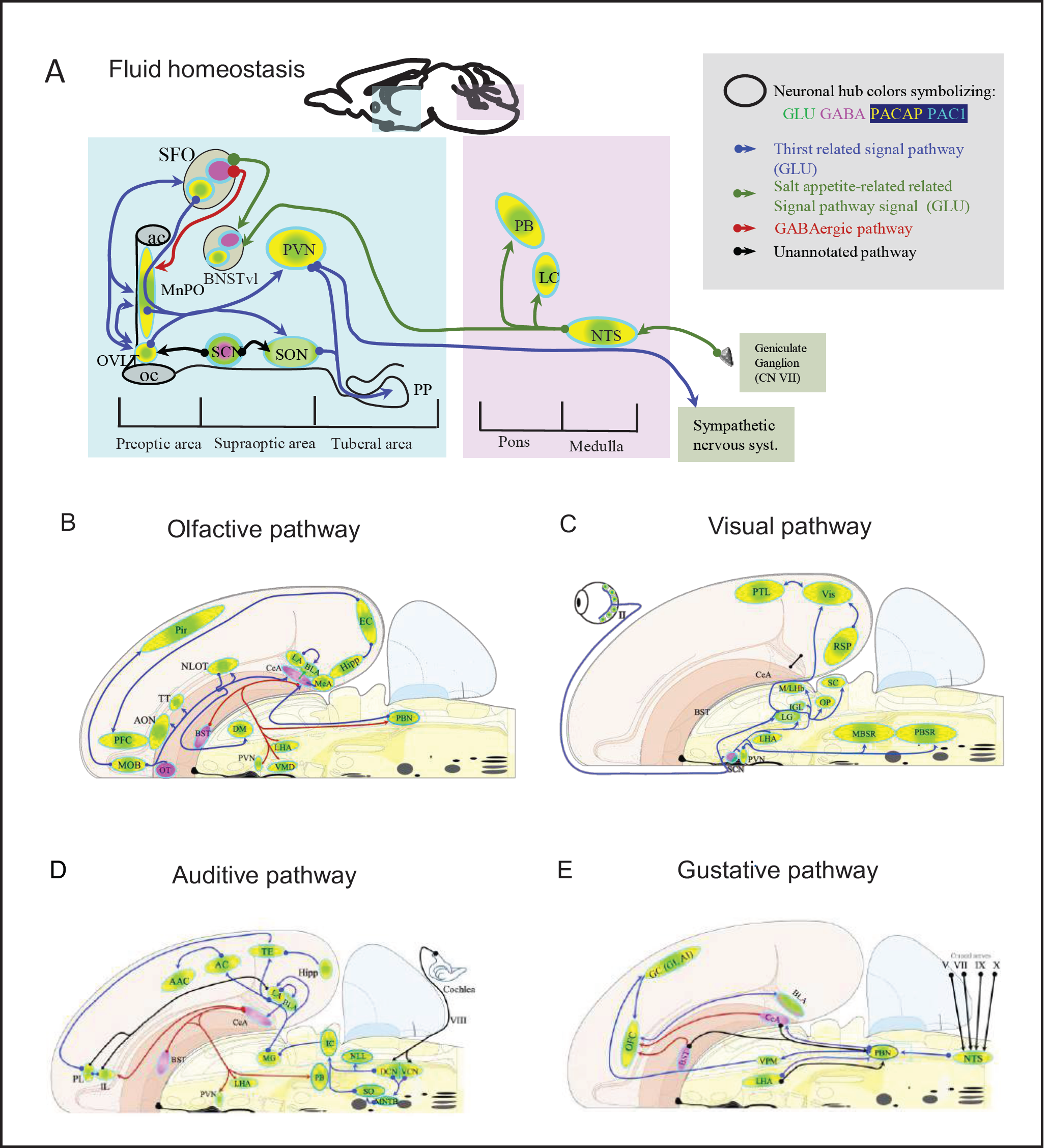
Mapping the spatial distribution of PACAP-PAC1 hubs within glutamate/GABA context in relevant sensory circuits in mice. **A**. Thirst and salt appetite-related pathways with PACAP-PAC1 glutamatergic / GABAergic signaling noted. The main figure is the enlargement of color-shaded areas of the box in the inset at the upper right, projected against a midsagittal section of mouse brain. Blue shaded area symbolizes the hypothalamus and pink shaded area the hindbrain. SFO, subfornical organ; MnPO, median preoptic nucleus; PVH, paraventricular nucleus; OVLT, organum vasculosum of the lamina terminalis; SON, supraoptic nucleus; SCN: suprachiasmatic nucleus (allostatic-anticipatory thirst, see main text); BNSTvl: bed nucleus of stria terminalis, ventrolateral division; PB: parabrachial complex; LC: locus coeruleus; NTS: nucleus of tractus solitarium; ac; anterior commissure; oc: optic chiasm; PP: posterior pituitary gland. **B**. Olfactory pathway. The projection neurons from the OB send their axons to the different structures of the olfactory cortex, among them the anterior olfactory nucleus (AON), the taenia tecta, the olfactory tubercle (OT), the piriform cortex (PC), the amygdaline complex (lateral amygdala LA, basolateral amygdala, BLA, central amygdala CeA, bed nucleus of stria terminalis BST) and entorhinal cortex (EC), and the nucleus of the lateral olfactory tract (nLOT). **C**. Visual pathway and circadian circuit for brain states. II: optic nerve; SCN: suprachiasmatic nucleus; PVN: paraventricular nucleus; LHA: lateral hypothalamic area; LG: lateral geniculate nuclei; IGL: intergeniculate leaflet; M/LHb: medial and lateral habenula; OP: olivary pretectal nucleus, SC: superior colliculus; Vis: visual area; PTL: parietal association area; RSP: retrosplenial area; MBSR: midbrain behavioral state related (pedunculopontine nucleus, substantia nigra, midbrain raphe nuclei; PBSR: pons behavioral state related (locus coeruleus, superior central nucleus of raphe, pontine reticular nucleus). **D**. Auditory pathway. VIII: cochlear nerve; DCN/VCN: dorsal&ventral cochlear nuclei; MNTB: medial nucleus of the trapezoid body; SO: superior olivary complex; NLL: Nucleus of the lateral lemniscus; IC: Inferior colliculus; MG: medial geniculate complex; AC: Auditory cortex; AAC: associate auditory cortex; TE: temporal association area; Hipp: hippocampus; LHA: lateral hypothalamic area; PVN: hypothalamic paraventricular nucleus; IL: infralimbic cortex; PL: prelimbic cortex. **E**. Gustative pathway. V, VII, IX, X: represent trigeminal, facial, glossopharyngeal & vagus nerves respectively. NTS: Nucleus of the solitary tract (nucleus of tractus solitarius); NA: nucleus ambiguus; PBN: Parabrachial nucleus; VPM: ventropostero medial nucleus of the thalamus; BLA: Basolateral amygdala; CeA: central Amygdala; BST: bed nucleus of the stria terminalis; LHA: lateral hypothalamic area; RF: reticular formation. ***Circuit diagrams used a modified version of Swanson’s flatmap*** http://larrywswanson.com/?page_id=1415 *and consulted https://sites.google.com/view/the-neurome-project/connections/cerebral-nuclei?authuser=0)*

**In the ventral (temporal pole) hippocampal formation**, we identified two cell populations with PACAP mRNA expression. One was the VGLUT1 mRNA-expressing mossy cells in the hilar region mentioned above, which were distributed from septo-dorsal to temporo-ventral hilus with increasing quantity. Mossy cells are major local circuit integrators and they exert modulation of the excitability of DG granule cells (Scharfman and Myers 2012, Sun, Grieco et al. 2017). The DG granule cells strongly expressed PAC1 (SI Fig. 3-E). Glutamatergic hilar mossy cells of the dentate gyrus can either excite or inhibit distant granule cells, depending on whether projections directly to granule cells or to local inhibitory interneurons (Scharfman and Myers 2012). However, the net effect of mossy cell loss on granule cell activity is not clear. Interestingly, dentate gyrus has a unique feature: there are two principal populations of glutamatergic cell type, the granule cells and the mossy cells. The former strongly expressed PAC1 and the latter strongly expressed PACAP, indicating that PACAP/PAC1 signaling may play a pivotal role for granule cell excitability.

A second population of PACAP mRNA-expressing neurons in **ventral hippocampus** was *a subset of CA3c pyramidal neurons*, previously photodocumented without comment in Figure 11-J of referenced report (Hannibal 2002). This represents a distinct and novel group of pyramidal neurons in ventral CA3 (Fig. 4, Panel A and C). These neurons strongly expressed PACAP mRNA, and co-expressed VGLUT1 (Fig. 4, panel B inset, panel D) and the calcium binding protein *calretinin* (Calb2) (Fig. 4 panel B), with the rest of pyramidal neurons expressing VGLUT1 but neither calretinin nor PACAP mRNA.

Retrohippocampal regions expressing PACAP and either VGLUT1 or VGLUT2 were: entorhinal area, prominently in the layer 5, parasubiculum, postsubiculum, presubiculum, and subiculum. This latter region, subiculum, together with the pyramidal layer of dorsal CA1, CA2 and CA3, exhibited large differences in PACAP mRNA expression strength between rat and mouse (SI Fig. 2). Developmental studies of these regions, as well as extended amygdala, may indicate a recapitulation of phylogeny by development that is relevant to the evolution of PACAP neurotransmission across mammalian species (Zhang and Eiden 2019).

#### 3.3 Cerebral cortex: structures derived from cortical subplate

The subplate is a largely transient cortical structure that contains some of the earliest generated neurons of the cerebral cortex and has important developmental functions to establish intra- and extra-cortical connections (Bruguier, Suarez et al. 2020). The concept of the subplate zone as a transient, dynamically changing and functional compartment arose from the combined application of functional and structural criteria and approaches (for a historical review see (Judas, Sedmak et al. 2010)). Here, we adapt our classification from that of the Allen Brain Map https://portal.brain-map.org/). Two noteworthy structures derived from the cortical subplate that expressed PACAP and PAC1 mRNAs are the claustrum (CLA) and the lateral amygdalar nucleus (LA).

##### 3. 3. 1 The claustrum (CLA)

Owing to its elongated shape and proximity to white matter structures, the claustrum (CLA) is an anatomically well-defined yet functionally poorly described structure, once speculated to be the “seat of consciousness” due to its extensive interconnections (Crick and Koch 2005). CLA is located between the insular cortex and the striatum: it is a thin sheet of gray matter considered as a major hub of widespread neocortical connections (Bruguier, Suarez et al. 2020). The CLA is recently reported to be required for optimal behavioral performance under high cognitive demand in the mouse (White, Mu et al. 2020). Consistent with recent work (White, Panicker et al. 2018), rat CLA receives a dense innervation from the anterior cingulate cortex (ACC), one of the most prominent PACAP mRNA-expressing regions in frontal cortex, co-expressing VGLUT1 and VGLUT2 mRNAs (Tab. 1 and Fig. 3, A-E & G) and is implicated in top-down attention (Zhang, Xu et al. 2016). The CLA interconnects the motor cortical areas in both hemispheres through corpus callosum (Smith and Alloway 2010), where PACAP-EGFP+ projections were reported (Condro, Matynia et al. 2016). Neuropeptides somatostatin (SOM), cholecystokinin (CCK) and vasoactive intestinal polypeptide (VIP) contents of the rat CLA have been reported (Eiden, Mezey et al. 1990). In mouse CLA, more than 80% of the neurons were VGLUT1 and VGLUT2 mRNA-co-expressing and less that 20% of the neurons expressed VGAT mRNA. PACAP content in rodent CLA has not been reported. In our study, we observed 10-15% of glutamatergic cells of the CLA co-expressed PACAP mRNA and almost 100% of cells expressed PAC1 mRNA.

##### 3. 3. 2 The cerebral cortex-derived components of endopiriform nucleus and amygdalar complex

The endopiriform nucleus and divisions of lateral, basolateral, basal medial and posterior amygdalar nuclei are, from a phylogenetic point of view, olfactory structures (Groor 1976), derived from the cerebral subplate whose main cell population is glutamatergic, co-expressing VGLUT1 and VGLUT2 (see Tab. 1). PACAP mRNA was strongly expressed in the lateral (dorsal) amygdala, anterior basomedial amygdala, posterior amygdalar nucleus, and with lower expression in the endopiriform nucleus and basomedial amygdala posterior subnucleus and sparse expression in the basolateral amygdala (SI Fig. 1, panel G).

#### 3.4 Structures derived from cerebral nuclei

The main structures expressing PACAP mRNA in striatum were the lateral septum (LS) and medial amygdala (MeA) (SI Fig. 1, panel G). Most of these neurons co-expressed both VGLUT1 and VGLUT2 mRNAs (see Tab. 1). In contrast to rat brain, where expression of PACAP mRNA is prominent in central amygdala and intercalated cells (IA), in the mouse PACAP mRNA-expressing cells were quite sparse in these structures (SI Fig. 2). PAC1 mRNA was intensely expressed in these mainly VGAT mRNA-expressing structures (Fig. 2 D-H, Tab. 2).

In structures within pallidum, PACAP mRNA was expressed, in order of abundance, in posterior, principal, stria extension and antero-medial divisions of BNST, and very sparsely, in the oval nucleus.

The strongest expression of PAC1 mRNA was observed on mainly GABAergic structures derived from cerebral nuclei. Figure 2-D-H show examples illustrating PAC1 mRNA co-expressing GABAergic (VGAT-expressing) neurons in some subcortical regions. More than 90% of cell population in all the nuclei of BNST co-expressed PAC1 mRNA (Fig. 2, D and F). In the oval subnucleus of BNST, we tested the three main GABAergic cell types that co-express somatostatin (Sst), parvalbumin (Pvalb) and corticotropin releasing hormone (Crh) and found all the three types of neurons co-expressed PAC1 mRNA (SI Fig. 3, panel I and I’, J and J’, K and K’). Table 2 summarizes the distribution, cell types and strength of main PAC1 expressing group in mouse cerebral nuclei.

**Table 2.**
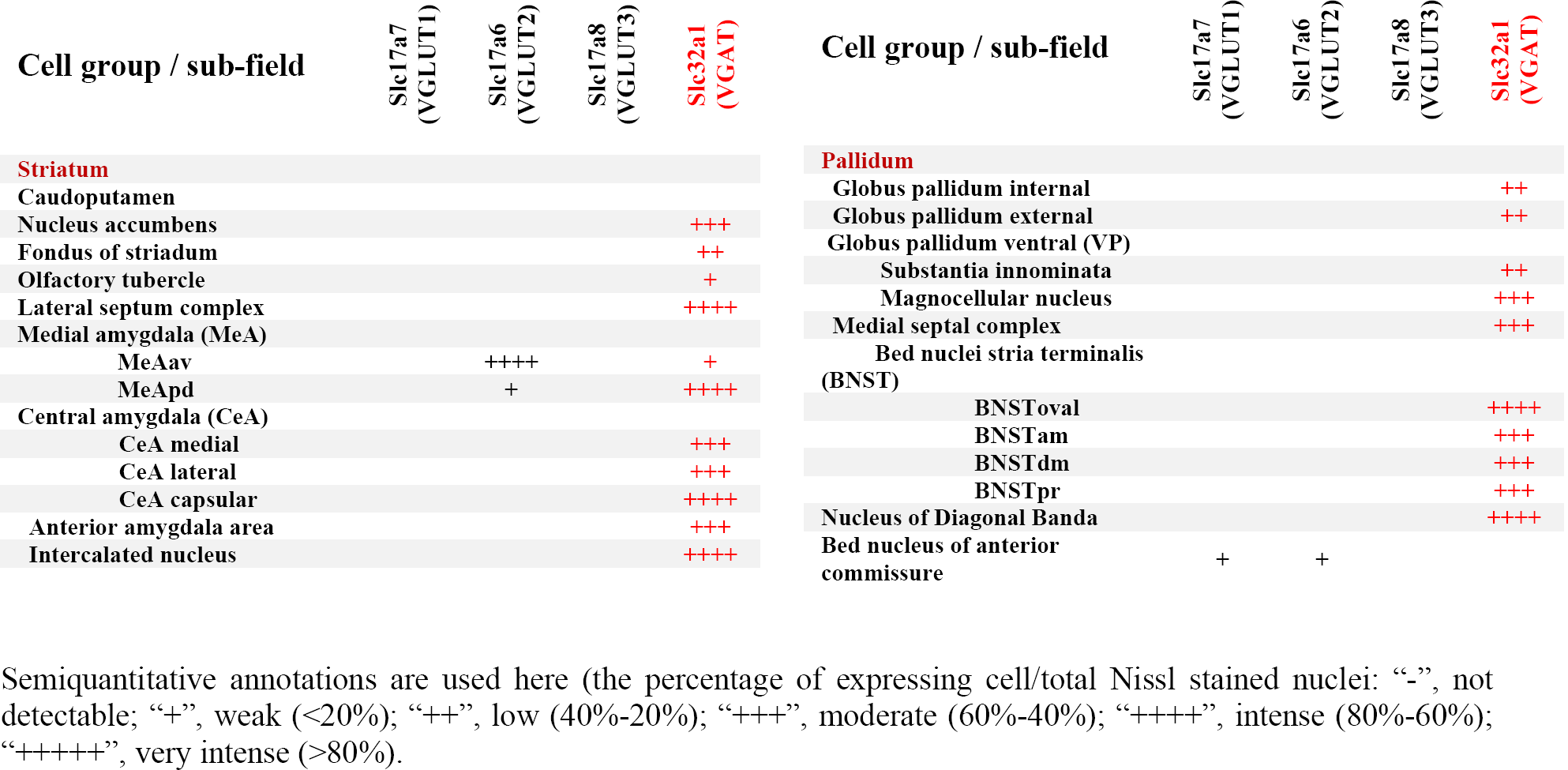
PAC1 expression and strength in striatum and pallidum structures

#### 3. 5 Brain stem

##### 3. 5. 1 Interbrain

###### 3. 5. 1. 1 Thalamus

PACAP mRNA is extensively expressed in thalamic nuclei (see Tab. 1), most prominently in medial geniculate complex, the nucleus reuniens, paraventricular thalamic nucleus (PVT) and the mediodorsal thalamic nucleus (MD) (SI Fig. J and N) neurons co-expressing Slc17a6 (VGLUT2). The medial geniculate nucleus or medial geniculate body is part of the auditory thalamus and represents the thalamic relay between the inferior colliculus and the auditory cortex. The nucleus reuniens receives afferent input mainly from limbic and limbic-associated structures, mediating interactions between the hippocampus and medial prefrontal cortex important for spatial working memory (Griffin 2015). It sends projections to the medial prefrontal cortex, the hippocampus, and the entorhinal cortex (Wouterlood, Saldana et al. 1990, McKenna and Vertes 2004), although there are sparse connections to many of the afferent structures as well. The prefrontal cortical-hippocampal connection allows regulation of neural traffic between these two regions with changes in attentiveness (Vertes, Hoover et al. 2007) as well as in resilience to stress (Kafetzopoulos, Kokras et al. 2018). All of the thalamic nuclei that express PACAP mRNA also express PAC1 mRNA (Tab. 1 and Fig. 3). The PVT and MD participate in many sensory information relays. In a recent study, their role in a key neural circuit for psychological threat-induced hyperthermia was reported (Kataoka, Shima et al. 2020). This circuit involves brain regions in the prefrontal pole, called the dorsal peduncular area (DPA 34a, also called dorsal taenia tecta, DTT, a main PACAP containing region mentioned in section 3. 2. 1, Fig. 3-B 34b) that senses social stress and mediates increased body temperature in response to it (Lin 2020). Neurons from the DP/DTT then project to and excite neurons in the dorsomedial hypothalamus (DMH, another PACAPergic nucleus in hypothalamus, Tab. 1, Fig. 3-B 52a, and *vide infra*), which in turn sends neuronal projections to the rostral medullary raphé (rMR, also a PACAP mRNA-expressing nucleus, Tab. 1, Fig. 3-A, 23, and description *vide infra*).

###### 3. 5. 1. 2 Epithalamus: habenula

Habenulae are bilateral triangular eminences of the stalk of the pineal gland, situated at the dorso-caudal end of the thalamus. Their medial divisions border the third ventricle. The habenula is considered as the relay hub where incoming signals from basal forebrain, for instance, diagonal band of Broca, lateral preoptic area, lateral hypothalamus, paraventricular nucleus, and entopeduncular nucleus, travel through the stria medullaris to habenula to be processed. The habenula then conveys the processed information to midbrain and hindbrain monoaminergic structures, such as ventral tegmental area, medial and dorsal raphe nuclei, and periaqueductal grey, through the fasciculus retroflexus. The habenula thus connects the cognitive-emotional basal forebrain to the modulatory monoaminergic area (Sutherland 1982). Medial habenula (MHb), was observed to express strongly PACAP mRNA in the dorsal half, in cells which co-express VGLUT1 (Slc17a7) or Slc17a6 VGLUT2 mRNAs (SI Fig. 1-H). In the lateral habenula, the PACAP-expressing neurons co-expressed VGLUT2 mRNA, and were mainly located in the central nuclei of the lateral habenula (SI Fig. 1-G), a region with rich input from hypothalamic peptidergic afferents including arginine vasopressin and orexin (Zhang, Hernandez et al. 2018). All those cells, both in lateral and medial habenula, co-expressed the calcium binding protein calretinin mRNA (Calb2).

###### 3. 5. 1. 3 Hypothalamus

Using the sensitive DISH method, a total of twenty-three hypothalamic nuclei were found to express PACAP mRNA (Tab. 1 and SI Fig. 1 and as well as in the website for public access https://gerfenc.biolucida.net/). Among the highest density PACAP mRNA-expressing cell clusters/nuclei (>80% of cells PACAP-positive) of hypothalamus are: organum vasculosum of lamina terminalis (SI Fig. 1-C), medial preoptic nucleus (Fig. 1, SI Fig. 1-C, D), subfornical organ (SI Fig. 1-F), ventro-medial hypothalamic nucleus (VMH, SI Fig. 1-G), subthalamic nucleus (SI Fig. 1-H), and lateral mammillary nucleus (SI Fig. 1-I). All these PACAP mRNA expressing cells co-expressed VGLUT2 mRNA. Other hypothalamic regions listed in Tab. 1 had lower density of expression and were also VGLUT2 co-expressing, such as paraventricular hypothalamic nucleus. VGAT/PACAP mRNAs co-expressing cells were sparsely distributed mainly in the anterior hypothalamic area (AHA), VMH, supramammillary and tuberomammillary nuclei (SI Fig. 1-O).

PAC1 mRNA expression in hypothalamus was extensive (Fig. 3 A-C) and in fact ubiquitous. The nuclei with highest PACAP mRNA expression mentioned above also had highest expression of PAC1 mRNA. In addition, the PVN, supraoptic nucleus (SON), suprachiasmatic nucleus (SCN), dorsomedial hypothalamic nucleus, arcuate hypothalamic nucleus (ARH), anterior hypothalamic nucleus, zona incerta, tuberal nucleus, lateral hypothalamic area, periventricular hypothalamic nucleus posterior, dorsal premammillary nucleus and supramammillary nucleus medial exhibited strong PAC1 expression. Both VGAT and VGLUT2 mRNA-expressing cells co-expressed PAC1 mRNA.

##### 3. 5. 2. Midbrain

We found moderate expression (between 40-60% of the cell population) of PACAP mRNA with VGLUT2 in the mainly sensory-related structures: inferior colliculus (IC), nucleus of the brachium of IC, midbrain trigeminal nucleus, and sparse expression (between 20-40%) in parabigeminal nucleus (Tab. 1, Fig. 3 A-D, SI Fig. 1 panels I-K and gerfenc.biolucida.net). We found strong co-expression of PACAP/VGLUT2 mRNAs (between 60-80% of the cell population) in motor related structures: superior colliculus, motor related subfield (SCm, SI Fig I and J) and periaqueductal gray (PAG, SI Fig I-K); moderate-sparse expression (between 20-60% of the cell population) in midbrain reticular nucleus, ventral tegmental area (VTA, SI Fig I and I9) and Edinger-Westphal nucleus. A PACAP population appeared which was VGLUT3-mRNA expressing (data not shown). These nuclei include: interfascicular nucleus of raphe, dorsal nucleus of raphe, central linear nucleus of raphe, rostral linear nucleus of raphe and olivary pretectal nucleus (Tab. 1). PAC1 mRNA was strongly expressed in the PAG and SC, although a moderate expression of this receptor was observed to be widespread in both glutamatergic (Slc17a6 and Slc17a8 expressing) and GABAergic (Slc32a1 expressing) cells (Fig. 3 A-D).

##### 3. 5. 2. Midbrain

We found moderate expression (between 40-60% of the cell population) of PACAP mRNA with VGLUT2 in the mainly sensorial related structures: inferior colliculus (IC), nucleus of the brachium of IC, midbrain trigeminal nucleus, and sparse expression (between 20-40%) in parabigeminal nucleus (Tab. 1, Fig. 3 A-D, SI Fig. 1 panels I-K and gerfenc.biolucida.net). We found strong co-expression of PACAP/VGLUT2 mRNAs (between 60-80% of the cell population) in motor related structures: superior colliculus, motor related subfield (SCm, SI Fig I and J) and periaqueductal gray (PAG, SI Fig I-K); moderate-sparse expression (between 20-60% of the cell population) in midbrain reticular nucleus, ventral tegmental area (VTA, SI Fig I and I9) and Edinger-Westphal nucleus. A PACAP population appeared which was VGLUT3-mRNA expressing (data not shown). These nuclei include: interfascicular nucleus of raphe, dorsal nucleus of raphe, central linear nucleus of raphe, rostral linear nucleus of raphe and olivary pretectal nucleus (Tab. 1). PAC1 mRNA was strongly expressed in the PAG and SC, although a moderate expression of this receptor was observed to be widespread in both glutamatergic (Slc17a6 and Slc17a8 expressing) and GABAergic (Slc32a1 expressing) cells (Fig. 3 A-D).

##### 3. 5. 3 Hindbrain

###### 3. 5. 3.1 Pons

In sensory-related structures, we found strongest PACAP mRNA expression (between 60-80% of the cell population) in the parabrachial complex (PBC), in all its subfields, although it was more intense toward its lateral divisions, external to the superior cerebellar peduncles (scp, Fig. 1-D and SI Fig. 1). The cells in those divisions, except the Koelliker-Fuse subnucleus (KF), were small cells mainly co-expressing VGLUT2 mRNA. In contrast, the PACAP cells in KF were bigger than the cells in the rest of the PBC divisions and strongly co-expressed Slc17a7 (VGLUT1). Other structures with moderate (20-40%) expression of PACAP mRNA and Slc17a7 (VGLUT1) were the lateral division of the superior olivary complex and the dorsal cochlear nucleus (SI Fig. M6). Lateral leminiscus nucleus and the principal sensorial nucleus of the trigeminal expressed PACAP mRNA sparsely in VGLUT2 mRNA expressing cells. In motor related structures, we found moderate PACAP mRNA co-expressed with VGLUT2 mRNA in the following structures: tegmental reticular nucleus, Barrington’s nucleus, dorsal tegmental nucleus, pontine grey (PG), pontine reticular nucleus, supratrigeminal nucleus and superior central nucleus raphe (Tab. 1 and Fig. 3-A-C for reference). In behavior state related structures, we found strong PACAP mRNA expression in locus coerulus co-expressing both VGLUT1 (some big cells) and VGLUT2 (in some small cells). The laterodorsal tegmental nucleus also expressed strongly the PACAP mRNA, co-expressing with VGLUT2 mRNA. Pontine reticular nucleus and superior central nucleus of raphe expressed PACAP mRNA with VGLUT2 mRNA in a less intense manner (Tab. 1). PAC1 mRNA was strongly expressed in pons structures, which also intensely expressed PACAP mRNA, such as PG, LC, PBC, DTN regions. Otherwise, the expression was observed to be widespread in both glutamatergic and GABAergic cell types (Fig. 3 A-D).

###### 3. 5. 3. 2 Medulla

In the medulla oblongata, PACAP mRNA was extensively expressed and generally in a sparse pattern. However, some of the nuclei showed strong-intense expression (60-80%): the nucleus of tractus solitarius (NTS), medial division (co-expressing mainly VGLUT1 mRNA), the NTS lateral division (co-expressing mainly VGLUT2), and the dorsal motor nucleus of the vagus nerve (DMX) (VGLUT1 mRNA expressing), the dorsal and ventral cochlear nuclei which co-expressed both VGLUT1 and VGLUT2 mRNAs. Details of other PACAP mRNA co-expressing VGLUT2 nuclei can be found in Tab. 1. The PAC1 mRNA expression in medulla is similar to pons, a widespread pattern with intense expression in NTS divisions, and other nuclei, which expressed PACAP mRNA (Fig. 3, A-F).

##### 3. 5. 4 Cerebellum

In the cerebellum, PACAP mRNA was expressed in all Purkinje cells, which co-expressed Slc32a1 (VGAT) (Fig. 1, 3 and SI Fig. 1-O). Figure 1, panels E-H show two cerebellar regions, paraflocculus (E and F) and central (G and H), low and high magnification respectively, where PACAP mRNA expression was higher than elsewhere. Purkinje cells, distributed in all regions of cerebellar cortex, are the are the most prominent population of GABA/PACAP co-expressing neurons of the brain. Some Golgi cells in the granule cell layer of paraflocculus and central regions also co-expressed PACAP and VGAT mRNAs (indicated with double pink arrowheads in Fig. 1 E-H). In these two cerebellar regions, some granule cells also expressed PACAP mRNA (indicated with single blue arrows, Fig. 1 E-H). In deep cerebellar nuclei, few PACAP mRNA-expressing cells were found in fastigial, interposed and dentate nuclei, with the former two co-expressing VGLU1 mRNA and the latter co-expressing VGLUT2. PAC1 mRNA expression was more limited than other brain regions analyzed above. It was mainly expressed in the VGAT mRNA-expressing Purkinje cells and sparsely expressed in VGLUT1 and VGLUT2 mRNA-expressing neurons in the deep cerebellar nuclei (Fig. 3, A-F).

### 4. PACAP→PAC1 signaling within sensory and behavioral circuits

Here, we analyze the chemo-anatomical aspects of PACAP/PAC1 mRNA expression using the results described above, but putting them into basic sensory circuit wiring maps, as well as in behavioral state and survival instinctive brain longitudinal structures, especially the hypothalamic hubs, based on existing classification schema (Swanson 2012, Sternson 2013, Swanson, Sporns et al. 2016, Zimmerman, Leib et al. 2017, Swanson 2018). Based on PACAP expression in these proposed sensory/behavioral circuits, we have addressed consequences of PACAP deficiency on neuronal activation and behavioral output in a mouse model of predator odor exposure and defensive behavior.

#### 4.1. PACAP-PAC1 co-expression in forebrain sensory system

##### 4.1.1. Thirst circuit for osmotic regulation

As shown above, PACAP mRNA was intensely expressed in all peri/para ventricular structures directly related to thirst and osmotic regulation (Fig. 7). These structures include SFO, OVLT, MnPO, and PVN (moderate expression) (SI Fig. 1 and Tab. 1). Other hypothalamic nuclei intrinsically related to osmotic control and anticipatory drinking are SON and SCN, which were strong PAC1-mRNA expressing.

**Figure 7.**
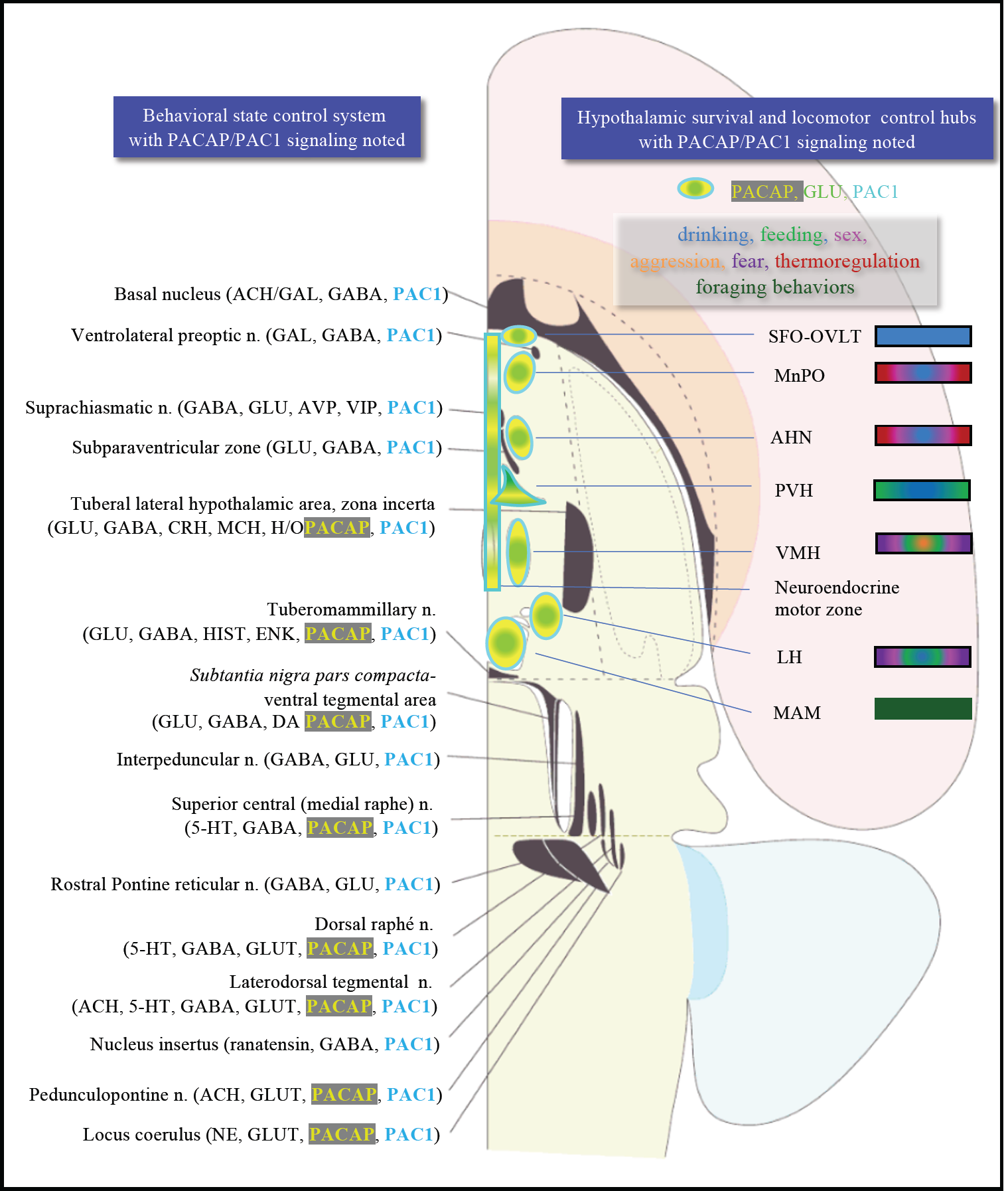
Presence of PACAP-PAC1 in major cell groups associated with behavioral state (left) and behavioral control (right column) control system and hypothalamic instinctive survival system. Left column: critical nodes for behavioral state symbolized by dark grey shaded objects, modified from in the longitudinal cell group-column of brain stem with key neurotransmitters annotated. ACH, acetylcholine; CRH, corticotropin-releasing hormone; DA, dopamine; ENK, encephalin: GABA, gamma-amino butyric acid; GAL, galanin; GLUT, glutamate; H/O, hypocretin/orexin; HIST, histamine; MCH, melanin-concentrating hormone; NE, norepinephrine; 5HT, serotonin. **Right column:** hypothalamic survival circuit that consisted of discrete hypothalamic regions contain interoceptors for a variety of substances and have neuronal afferences from primary sensory systems to control the secretory and instinctive motor outputs. The rectangle in the midline represents the neuroendocrine motor zone for secretion of hypophysiotropic hormones, which include thyrotropin-releasing hormone, corticotropin-releasing hormone, growth hormone-releasing hormone, somatostatin, gonadotropin-releasing hormone, dopamine, neurotensin. SFO, subfornical organ; OVLT, organum vasculosum of lamina terminalis; MnPO, preoptic nucleus; AHN, anterior hypothalamic area; PVH, paraventricular hypothalamic nucleus; VMH, ventromedial hypothalamic nucleus; LH, lateral hypothalamic area; MAM, mammillary nucleus (for general reference see (Swanson 2012)).

**Figure 8.**
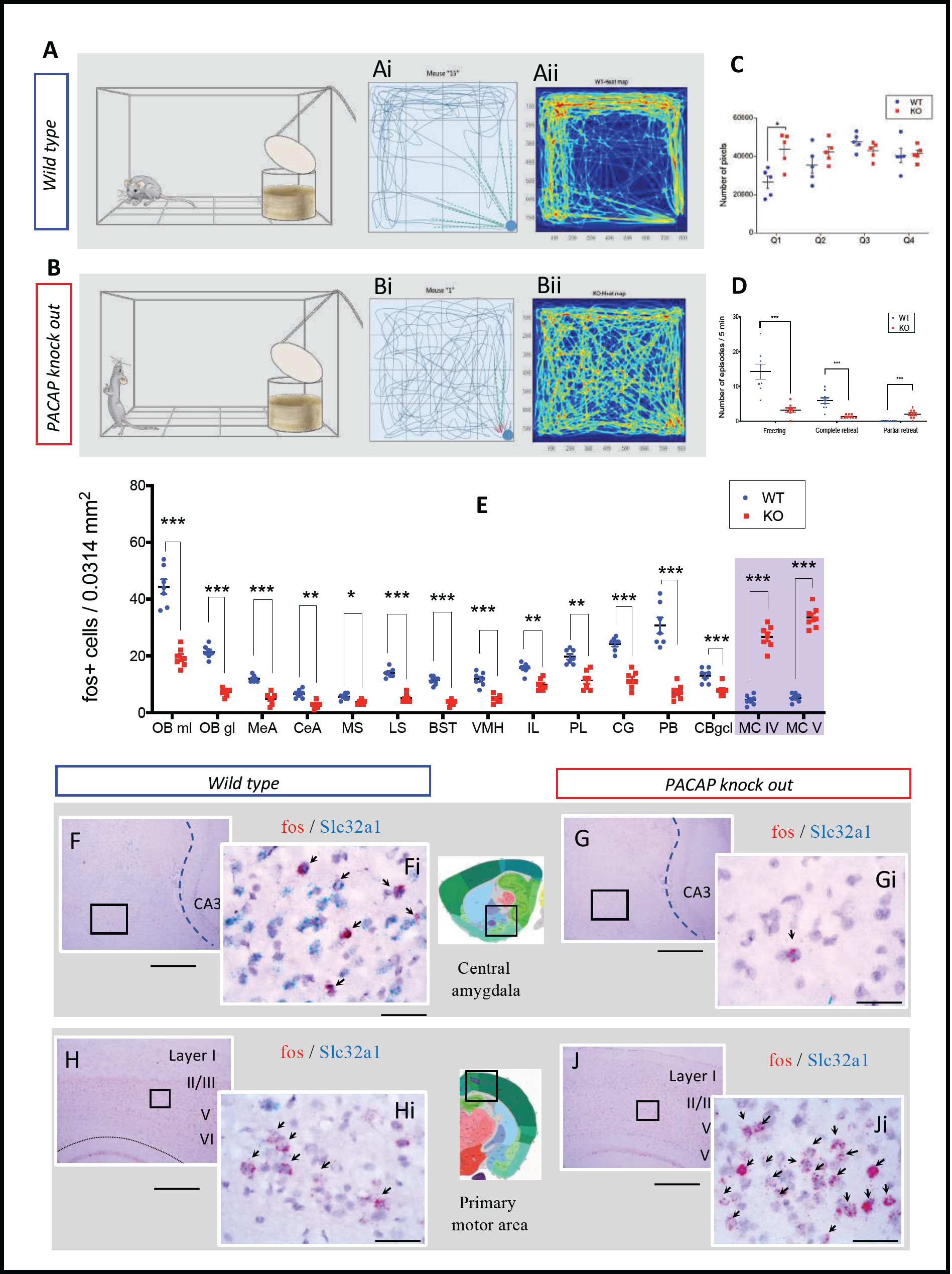
PACAP deficient mice exhibited aberrant predator odor response and defensive behavior. A and B: Schematics of behavioral procedure using a modified OFT to study defensive behavior during predator odor exposure, with typical behaviors of wild type (A: freezing) and PACAP KO (B, wandering, hyperactive and jumping) symbolized. Ai and Bi: representative 2D movement tracking in the OFT of WT (Ai) and OK (Bi) mice, with dashed lines symbolizing complete (green lines) /incomplete (red lines) approach/sniff > retreat cycles. Aii and Bii: 2D movement heat-map WT, *n*=9 mice; KO, *n*=9 mice C: pixel analysis reflecting total distance traveled in each of the four quadrants, being the Q1 the container located quadrant (clockwise numbering). The number of pixels per quadrant for mice in the KO group was compared with mice in the WT group following one-way ANOVA. The difference was significant in quadrant 1. WT average = 26500 and KO average = 43640, *p<0.05 (t= 3.35, df= 7.76, F=1.42). D: Number of freezing, complete sniff>retreats or partial sniff>retreats behaviors, assessment was made every five seconds during a 5 min test period. Student t-test, showed in KO mice a significant decrease (*** *p<0.001)* in freezing (WT:14.29 ±2.18 vs KO: 3.18 ± 0.64) and complete sniff>retreat behaviors freezing (WT: 5.89 ± 0.84 vs KO: 1.33 ± 0.17) and an increase in the partial sniff>retreat behaviors (WT: 0 vs KO: 2 ± 0.41). H. Number of cells expressing fos RNA 45 min after the predator odor test were quantified in a 0.0314 mm^2^ area and statistic significant differences between KO and WT were determined using a multiple t-test and the Bonferroni-Dunn method. There was a significant decrease in the expression of *fos* in the following regions of KO mice: olfactory bulb mitral layer “OB ml” (WT: 44.4 ± 2.60 vs KO: 19.43 ±1.25); olfactory bulb granule layer “OB gl”(WT: 21.43 ±0.84 vs KO: 7.29 ± 0.52); medial amygdala “MeA” (WT: 12 ± 0.44 vs KO: 5 ± 0.76); central amygdala “CeA” (WT: 6.71 ± 0.57 vs KO: 2.71 ± 0.47); medial septum “MS” (WT: 5.71 ± 0.42 vs KO: 3.71 ± 0.29); lateral septum “LS” (WT: 14 ± 0.62 vs KO: 5.14 ± 0.56); bed nucleus of stria terminalis “BST” (WT: 11.43 ± 0.57 vs KO: 3.57 ± 0.37); ventromedial hypothalamus “VMH” (WT: 11.86 ± 0.91 vs KO: 4.86 ± 0.50); infralimbic cortex “IL” (WT: 15.71 ± 0.714 vs KO: 9.86 ± 0.74); prelimbic cortex “PL” (WT: 19.71 ± 0.97 vs KO: 11.43 ± 1.23); cingulate cortex “CG” (WT: 24.28 ±0.92 vs KO: 11.57± 1.15) and parabrachial nucleus “PB” (WT: 30.71 ± 2.82 vs KO: 7.29 ± 1.02) and granule cell layer of cerebellum “CBgcl” (WT: 13.14 ± 0.96 vs KO: 8 ± 0.76). In contrast, the number of *fos* nuclei was augmented in two motor related areas in KO animals related to WT: motor cortex, layer IV “MC IV” (WT: 4.43 ± 0.65 vs KO: 26.71 ± 1.54) and motor cortex, layer V (WT: 5.28 ± 0.57 vs KO: 33.57 ± 1.43). F and G: example of reduced *fos* expression pattern in central amygdala in KO mice and H and J augmented *fos* expression in primary motor cortex of KO mice. Scale bars: 200 µm for low magnification and 20 µm for high magnification insets. Statistic significant differences are depicted as *** *p<0.001, ** p<0.01*, * *p<0.05*.

The SFO is an embryonic differentiation of the forebrain roof plate, in a dorsal region between the diencephalon (interbrain, thalamus) and the telencephalon (endbrain). This nucleus lacks a normal blood-brain barrier, and so its neurons are exposed directly to peptide hormones in the blood. One such hormone is angiotensin II, whose blood levels are elevated upon loss of body fluid due to dehydration or hemorrhage. Hence, the SFO is a *humorosensory organ* that detects hormone levels in the circulation to control drinking behavior and body water homeostasis. The SFO is situated immediately dorsal to the third ventricle and contains intermingled populations of glutamatergic (VGLUT2-PACAP-PAC1) and GABAergic (VGAT-PAC1) neurons with opposing effects on drinking behavior. Optogenetic activation of SFO-GLUT neurons stimulates intensive drinking in hydrated mice, whereas optogenetic silencing of SFO-GLUT neurons suppresses drinking in dehydrated mice (Zimmerman, Leib et al. 2017). By contrast, optogenetic activation of SFO-GABA neurons suppresses drinking in dehydrated mice (Bichet 2018). SFO-GLUT projections to the median preoptic nucleus (MnPO) and OVLT drive thirst, whereas SFO-GLUT projections to the ventrolateral part of the bed nucleus of the stria terminalis (BNSTvl) promote sodium consumption (Zimmerman, Leib et al. 2017). SFO-GLUT projections to the paraventricular (PVN) and supraoptic (SON) nuclei of the hypothalamus have not yet been functionally annotated with cell-type specificity, but classic models suggest that these projections mediate secretion of arginine vasopressin (AVP) and, in rodents, oxytocin (OXT) into the circulation by posterior pituitary (PP)-projecting magnocellular neurosecretory cells (MNNs). Recent studies also demonstrated that these MNNs possess ascending projections innervating limbic structures such as amygdala, hippocampus, lateral habenula and lateral hypothalamus (Hernandez, Vazquez-Juarez et al. 2015, Hernandez, Hernandez et al. 2016), in a cell-type specific manner (Zhang and Hernandez 2013, Zhang, Hernandez et al. 2016, Zhang, Hernandez et al. 2018). When these are activated, their central collaterals can exert motivational effect on exploration and drinking behavior.

Thirst and AVP release are regulated not only by the classical homeostatic, interosensory plasma osmolality negative feedback (through SFO as a humorosensory organ), but also by novel, exterosensory, anticipatory signals (Gizowski, Zaelzer et al. 2016). These anticipatory signals for thirst and vasopressin release converge on the same homeostatic neurons of circumventricular organs that monitor the composition of the blood. Acid-sensing taste receptor cells (which express polycystic kidney disease 2-like 1 protein) on the tongue that were previously suggested as the sour taste sensors also mediate taste responses to water. Recent findings obtained in humans using blood oxygen level-dependent (BOLD) signals demonstrating that the increase in the lamina terminalis (LT) BOLD signal observed during an infusion of hypertonic saline is rapidly decreased after water intake well before any water absorption in blood. This is relevant in the context of this paper since the MnPO of the hypothalamus has been shown to mediate this interesting phenomenon, integrating multiple thirst-generating stimuli (Allen, DeNardo et al. 2017, Gizowski and Bourque 2017), however, functionally annotated cell-type specific circuitry has not been clarified. The highest expression level of PACAP mRNA was observed in MnPO. Together, these observations open new possibilities to further understand the role of PACAP-PAC1 signaling within this nucleus for homeostatic and allostatic control.

Information about plasma sodium concentration enters the circuit through specialized aldosterone-sensitive neurons in the NTS, a strongly PACAP-expressing nucleus in the medulla, that expresses 11β-hydroxysteroid dehydrogenase type 2 (NTS-HSD2 neurons) (Zimmerman, Leib et al. 2017, Bichet 2018), which promote salt appetite and project to the LC (PACAP-expressing), PBN (intensely PACAP-expressing) and BNSTvl (PACAP was moderately and PAC1 was strongly expressed) (Fig. 6-A).

##### 4.1.2 Olfactory pathway

In the main olfactory system, input from olfactory sensory neurons reaches the main olfactory bulb (MOB), and the axons of the projection neurons in MOB travel to the anterior olfactory nucleus (AON), piriform cortex (Pir), and amygdala for additional processing (Wacker, Engelmann et al. 2011). This multi-step pathway brings olfactory information to be processed in multiple areas of the cerebral cortex, including the anterior AON and Pir. The AON, a cortical area adjacent to the olfactory bulb, is part of the main olfactory pathway. A parallel system, the accessory olfactory system, brings information, for instance that conveyed by pheromones, from the vomeronasal organ into the accessory olfactory bulb, which innervates the MA, BNST, and cortical amygdala (Fig. 10). Although it was originally assumed that only the accessory olfactory system processed pheromonal and other socially relevant odors, more recent evidence suggests that social information is processed by both pathways (Wacker and Ludwig 2012).

**Figure 9.**
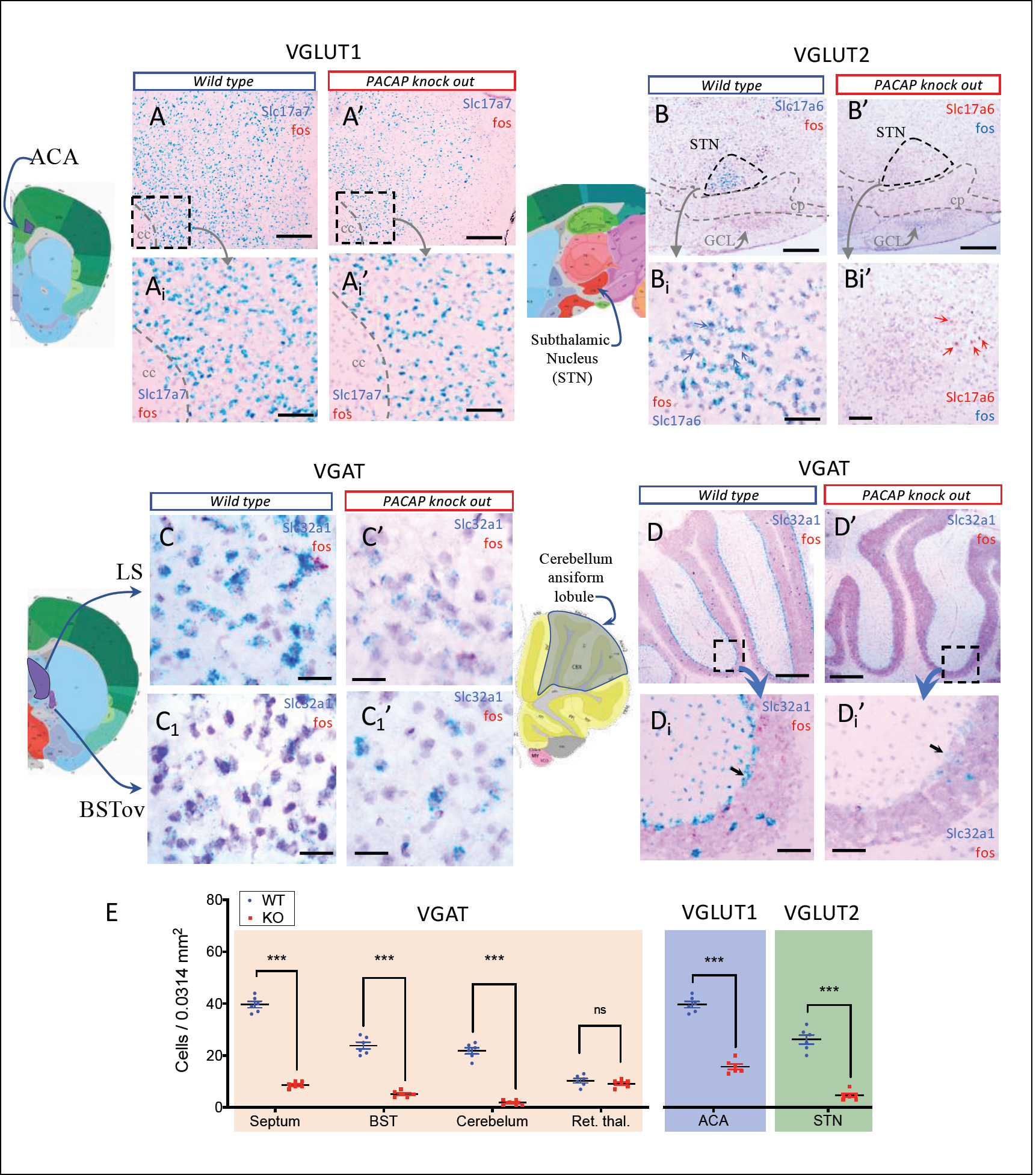
PACAP deficient (KO) mice showed significant down-regulation of vesicular transporters for glutamate and GABA in regions where Adcyap1 or Adcyap1r1 were strongly expressed in WT mice. A-D: examples of in situ hybridization using RNAscope method showing down-regulation in KO mice (A’, B’, C’, D’), of Slc17a7 (VGLUT1) in anterior cingulate area (ACA) (panels As), Slc17a6 (VGLUT2) in hypothalamic subtalamic nucleus (STN) (Panels B) and of Slc32a1 (VGAT) in lateral septum (LS) and bed nucleus of stria terminalis, oval subnucleus (BSTov) (panels Cs), and cerebellar cortex, the ansiform lobule’s, Purkinje’s cells (panels Ds). Note that the feature of reduced expression of Slc32a1 at both single cell and cell density levels, can be clearly observed (arrows) in the Purkinje cells. Number of cells expressing VGAT RNA (orange shading), VGLUT1 RNA (blue shading) and VGLUT2 (green shading) were quantified in a 0.0314 mm^2^ area and statistically significant differences between KO and WT were determined for septum (WT: 39.67 ± 1.23 vs KO: 8.67 ± 0.49); BST (WT = 23.83 ± 1.35 vs KO = 5.17 ± 0.48); ansiform lobule of cerebellum (WT: 21.83 ± 1.19 vs KO: 1.83 ± 0.4); anterior Cingulate Area (WT: 39.67 ± 1.23 vs KO: 15.67 ± 1.05) and subthalamic nucleus (WT: 26.17 ± 1.78 vs KO: 4.67 ± 0.76). Finally, we chose the reticular thalamic nu. (WT: 10.33 ± 0.88 vs KO: 9.17 ± 0.6), a region that does not containg PACAP expressing cells, as our negative control. Statistic differences are depicted as *** *p<0.001 and* ns: not significant). Scale bars: A & A’: 300 µm; Ai & Ai’: 100 µm; Bs and Cs: 20 µm; D & D’: 500 µm; D1 & D1’: 100 µm.

**Fig. 10.**
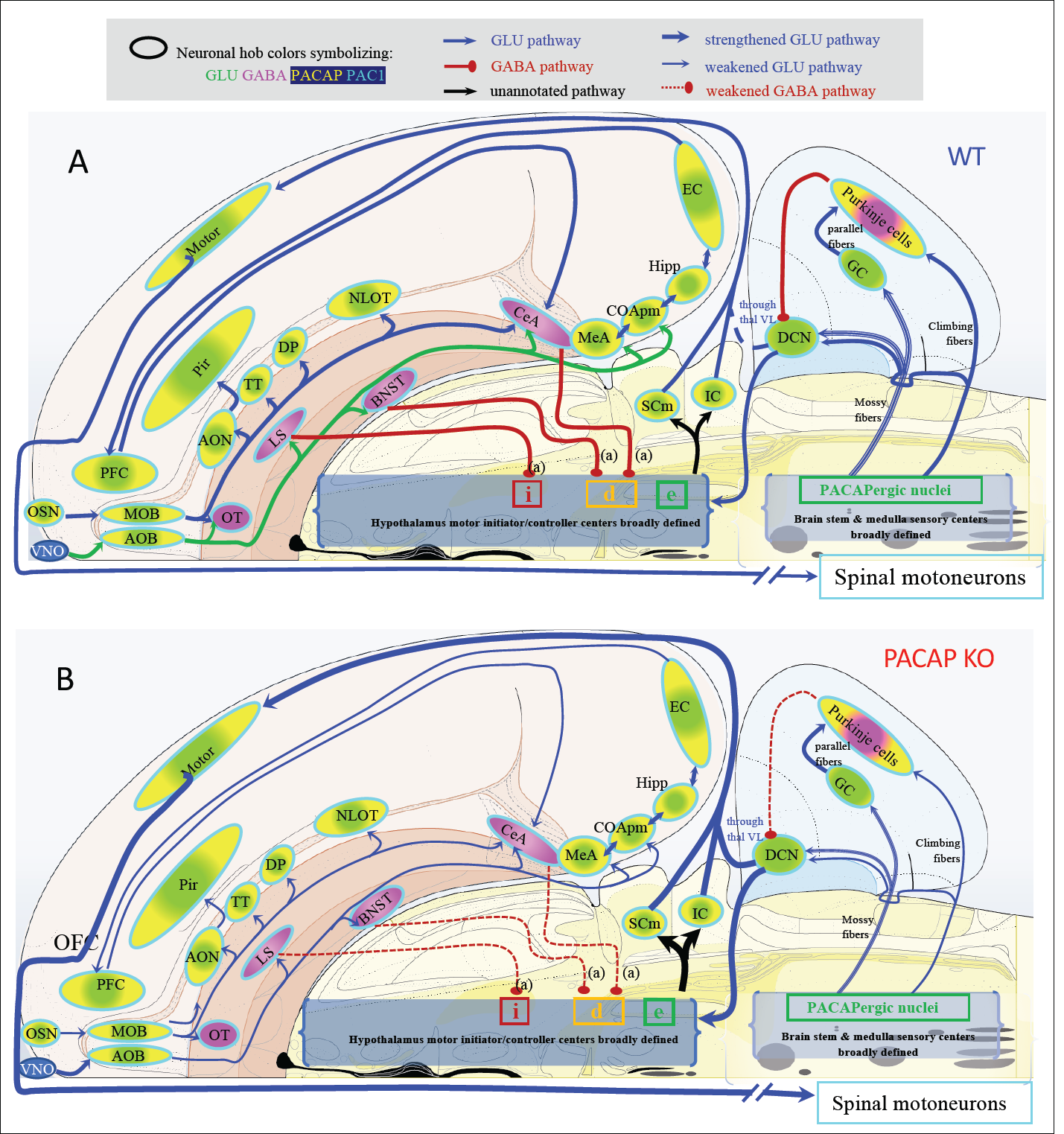
Proposed model to interpret how PACAP deficiency influence the olfactory information salience processing to impact motor output, bases on the analysis of this study. (A) Schematic representations of the mouse peripheral and central olfactory pathways to cognitive centers and motor higher control center in brain stem under normal condition. PACAP-Pac1 glutamatergic/GABAergic signaling is noted by colors. B: PACAP deficiency weakens the corresponding neuronal hubs inducing ultimately unbalanced excitation/inhibition in sensory-cognitive and motor pattern initiator and controller center, which may underlie the hyperactivity and attention deficit to salient olfactory stimulus during predator odor exposure test. OSN: olfactory sensorial neurons; VNO: vomeronasal organ; MOB: main olfactory bulb; AOB: accessory olfactory bulb; AON, anterior olfactory n.; TT: taenia tecta; OT: olfactory tubercle; DP: dorsal peduncular area; PC: piriform cortex; NLOT: n. lateral olfactory tract; EC: entorhinal cortex; PFC: prefrontal cortex; COApm: corticoamygdalar, posteromedial; MeA and CeA: medial and central amygdala; LS: lateral septum; BNST: bed nucleus of stria terminalis; SCm: superior colliculus, motor related; IC: inferior colliculus; DCN: deep cerebellar nuclei; GC: granule cells; thal VL: thalamic ventrolateral n.; i: inhibition; d: disinhibition; e: excitation.

The olfactory system appears especially to use PACAP/PAC1 as one of its main modes of co-transmission (see section 3. 2. 1). The above two olfactory circuits are interconnected with diencephalon and hindbrain structures prominently for reproductive and defensive behaviors, such as VMH, lateral hypothalamic areas, PVN and PBC in the pons, which are PACAP/PAC1 and VGAT mRNAs-co-expressing cell populations.

##### 4. 1. 3 Visual pathway and the circadian circuits for brain states

There is an enormous literature on the visual system and we only touch on selected hubs here to emphasize the PACAP-PAC1 signaling role for visual information processing (Fig. 6-C). PACAP’s role in the visual system was first discovered via identification of the PACAP immunopositive retinal ganglion cells (RGC), soon after the discovery of PACAP itself (Hannibal, Ding et al. 1997), which project to the SCN. Here, using DISH method, we report Adcyap1 co-expressed with Slc17a6 (VGLUT2) and retina ganglion cells. PACAP mRNA expression intensity followed a circadian oscillation mode (see section 3.1). The RGC sends visual information through its axons forming the optic nerve, chiasm and optical tract and accessory optical tract, with its first offshoot to SCN for brain state modulation. The SCN was found intensely expressing PAC1 in both VGAT and VGLUT2 mRNA-expressing neurons widely and homogenously distributed, at variance with the observation in rat that PAC1 is expressed mainly in the ventral part of the SCN (Hannibal, Ding et al. 1997). Leaving the optic chiasm, the optic tract courses latero-caudally and emits a second offshoot, the accessory optical tract, splits off and courses to the midbrain, where it ends in three terminal nuclei, i.e. medial, lateral and dorsal terminal nuclei. They play an important role in controlling eye movements and are thus parts of the motor system and were all PACAP mRNA-expressing. The main optic tract continues on to end in the superior colliculus (SC) of the midbrain, after giving off collaterals to the lateral geniculate complex (LG) of the thalamus and to the olivary pretectal nucleus (OP). The dorsal part of the LG then projects to primary visual cortex (Vis), whereas the OP is involved in visual reflex, and the superior colliculus has two main roles: projecting to the motor system, and projecting to secondary visual cortical area via thalamus (Swanson 2012). All above optic nerve/tract/accessory tract targeted structures were observed to have intense expression of PAC1 mRNA. The visual cortex, parietal associated area, and retrosplenial area strongly expressed PACAP mRNA (Tab. 1 and SI Fig. 1, J, K).

PAC1 mRNA was strongly expressed in SCN (Fig. 6-C and Fig. 7), which projects in turn to several PAC1-expressing regions, which control brain state (sleep-wake cycle), such as midbrain behavioral state related structures (MBSR: pendunculopontine, midbrain raphe nuclei, all PACAP mRNA expressing) and pons behavioral state related (PBSR: locus coeruleus, superior central nucleus of raphe, pontine reticular nucleus, all PACAP-mRNA expressing).

#### 4.2 PACAP-PAC1 co-expression in the central processing of ganglion cell sensory systems

The sensory ganglion cells were observed to express PACAP shortly after its discovery (Sundler, Ekblad et al. 1996). PACAP mRNA and peptide levels are increased in sensory ganglion cells within one day following axotomy, suggesting possible roles in neuronal protection, differentiation, nerve fiber outgrowth, and/or restoration of perineuronal tissue upon neuronal damage. The sensory ganglion cells send their axons to primary sensory nuclei in the dorsal medulla, which include: a) auditory system, which ends in the cochlear nuclei (Fig. 6-E); b) vestibular system, which ends in the vestibular nuclei (not included in this analysis); c) gustatory system which ends in the rostral nucleus of *tractus solitarius* (NTS, Fig. 6-E); and d) vagal/glossopharyngeal visceroceptive system, which ends in the caudal nucleus of the NTS (not included in this analysis). The special sensory nuclei in the medulla are all derived in the embryo from a highly differentiated, dorsal region of the primary hindbrain vesicle, the rombic lip (Fig. 5.14 of (Swanson 2012)) – they were all observed to co-express PACAP and VGLUT2 mRNAs from very intense (NTS) to moderate levels (DCN and VCN) (see Tab. 1, Fi. 2 A-E, SI Fig. 1 and website).

Here we analyze the PACAP/PAC1 participation in the auditory and gustatory central processing circuits. 20

##### 4.2.1. Auditory pathway

Auditory information, for example the spectrum, timing, and location of sound is analyzed in parallel in the lower brainstem nuclei, i.e., dorsal and ventral cochlear nuclei (DCN and VCN), medial nucleus of trapezoid body (MNTB), superior olivary complex (SO), and nuclei of the lateral lemniscus (NLL). These hindbrain structures (NLL and SO) project to the inferior colliculus (IC, in midbrain) through both the excitatory and inhibitory inputs to IC. From IC the information travels toward thalamus via relay in medial geniculate complex (MG), to reach the auditory cortex, associate auditory cortex, temporal association area; hippocampus, which subsequently projects to prefrontal cortices (Ono and Ito 2015). Main neuronal structures that are relevant to the auditory pathway are shown in the schematic diagram of Fig. 6-B. The glutamatergic population of these structures were all found to co-express PACAP, VGLUT2 and/or VGLUT1, as well as PAC1 mRNAs.

The mainly VGAT mRNA-expressing structures located in subcortical regions which strongly co-expressed PAC1 mRNAs (Fig. 2 D-G and Tab. 2), include, for instance, BST and the CeA with its three subdivisions (Fig. 6-D, structures symbolized in pink-GABAergic, and blue outline–PAC expressing) participate in cognitive and emotional auditory information, obtaining auditory sensory input from brain stem>midbrain>thalamic MG, which project to lateral amygdala (LA, a main PACAPergic nucleus derived from cortical plate) which send input to the cognitive centers through basal lateral amygdala (BLA, Fig. 2, F and G).

##### 4.2.2. Gustatory pathway

Chemicals (taste substances) in foods detected by sensory cells in taste buds distributed in the oropharyngeal epithelia are recognized as tastes in the gustatory cortex (GC), including granular insular cortex (GI) and agranular insular cortex (AI), also called dysgranular insular cortex (DIC), which is the primary gustatory cortex (Fig. 6-E). Between the peripheral sensory tissue and the GC, many neuronal hubs participate in the gustatory information processing, beginning with the geniculate ganglion (GG), of the facial nerve, cranial nerve CN-VII which receive taste stimuli from the anterior 2/3 of the tongue; the petrosal ganglion (PG), of the glossopharyngeal nerve, CN-XI, which receive the taste stimuli from the posterior third of the tongue; and nodose ganglion (NG) ganglia, also called the inferior ganglion of the vagus nerve (CN-X, which innervate the taste buds on the epiglottis). The lingual nerve, a branch of mandibular nerve, from trigeminal nerve (CN-V), also innervates the anterior 2/3 portion of tongue. All of these ganglia express PACAP as their co-transmitter and this expression is potentiated during injury (Sundler, Ekblad et al. 1996). Taste information from sensory ganglia of CN V, VII, IX, X, project to the rostral NTS of the medulla, which intensely expressed PACAP and PAC1 mRNAs. NTS neurons send taste information to parabrachial nucleus (PB, also intensely expressed PACAP and PAC1 mRNAs) and then to the parvocellular division of ventral posteromedial nucleus VPM of the thalamus, which project to GC, as well as to the cognitive centers, central amygdala (CeA), bed nucleus of stria terminalis (BNST) and lateral hypothalamus (LH) that these cognitive centers also reciprocally innervate the PB (Halsell 1992). In Fig. 6-E, the basic wiring of taste circuit was based on the literature (Halsell 1992, Carleton, Accolla et al. 2010), and modified with PACAP-PAC1 signaling annotated.

#### 4. 3 PACAP-PAC1 signaling in major cell groups associated with behavioral state control system and hypothalamic instinctive survival system

In the course of a day, brain states fluctuate, from conscious awake information-acquiring states to sleep states, during which previously acquired information is further processed and stored as memories (Tukker, Beed et al. 2020). Anatomically and chemically distinct neuronal cell groups stretching from medulla, pons, midbrain, through the hypothalamus to the cerebral nuclei all participate in modulation of behavioral state (Swanson 2012). We have briefly referred to sleep and waking behavioral states in analysis of the hypothalamic SCN in the visual/photoceptive pathway that project to midbrain and hindbrain behavioral state structures broadly defined. In the Fig. 7, left column we complemented the model of behavioral state, cell groups and chemical signatures previously presented (Swanson 2012), adding PACAP-PAC1 as informed by the current study. Three behavioral state-related glutamatergic structures in medulla, *i.e. raphe magnus, raphe pallidus* and *raphé obscurus* are not represented in the left column because of recognizing the original figure design of the author (Fig. 9.5, (Swanson 2012)).

In the right column of Fig. 7, we present the main hypothalamic survival and locomotor control hubs with PACAP/PAC1 signaling noted. These hubs are SFO, OVLT, MnPO, anterior hypothalamic nucleus (AHN), PVN, VMH, lateral hypothalamus (LH), mammillary nucleus (MAM), and periventricular neuroendocrine motor zone. Recent evidence suggests that these cells groups control the expression of motivated or goal-oriented behaviors, such as drinking, feeding, sex, aggression, fear, foraging behaviors (Fig. 7, color-filled rectangles represent the correlative behaviors with anatomical structure based on literature (Sternson 2013)). The rostral segment of this behavior control column has controllers for the basic classes of goal-oriented ingestive, reproductive and defensive behaviors, common to all animals, where the caudal segment has the controllers for exploratory behavior used to obtain any goal object (Swanson 2012).

Other relevant structures more caudal in this longitudinal brain stem column are the reticular part of the subtantia nigra (not labeled in Fig. 7) which is involved in the control of orientating movements of the eyes and head via projecting to the superior colliculus, the ventral tegmental area (VTA), PACAP mRNA expressing, which together with nucleus accumbens (PAC1 mRNA strongly expressing) and subtalamic nucleus (PACAP strongly expressing), form the hypothalamic locomotor region.

After analyzing the above presence of PACAP>PAC1 in sensory-cognitive to motor output, a question emerged: what would be the behavioral and neuronal activation consequences if PACAP signaling were deficient?

#### 4.4. PACAP knockout impairs predator odor salience processing via reducing neuronal activation and vesicular transporter expression in key PACAP-PAC1 nuclei

To assess the behavioral implications of PACAP action within one of these circuits, we examined the effect of knockout of PACAP expression in brain on defensive behavior initiated via olfaction, using a predator odor paradigm (Fig. 8, A-B). We employed a modified open-field box with a lidded containerin which cat urine-saturated litter was placed. Using wild-type (WT) and PACAP-deficient (KO) C57Bl/6N mice (Hamelink, Tjurmina et al. 2002), we assessed the defensive locomotion patterns during cat odor exposure, which include displacement pattern in the four quadrants of the box; approach to the stimulus (urine-containing cat litter) with complete retreats and incomplete retreats; and periods of immobility (freezing behavior). DISH for fos expression and Slc17a7, Slc17a6 and Slc32a1 (VGLUT1, VGLUT2 and VGAT) expression in the two experimental groups was performed and analyzed *post hoc*.

Cat urine triggered purposeful movement such as sniff>retreat in WT subjects (Fig. 8, Ai, green dashed lines indicate complete sniff>retreat cycles), contrasting the movement patterns exhibited by KO subjects that most of the sniff>retreat cycles were short distances and the mice often returned to the container location area (Fig. 8, Bi, red dashed lines indicate incomplete/short sniff>retreat cycles). PACAP-deficient mice had significant reduction of complete retreats (p<0.001) and significant augmentation of incomplete retreats (p<0.001) (Fig. 8-D). Figures 8 Aii and Bii show the X-Y dimension movement heat-maps for each group. Movement analysis by quadrants showed a significant increase of pixel values of the KO group in the quadrant where the odorant container was located (Fig. 8-C, p<0.05, C1). It is worth mentioning that movement in Z dimension (repetitive jumping), characteristic of PACAP-deficient mice (Hashimoto, Shintani et al. 2001) was not represented in the X-Y dimension heat maps and investigating the phenomenon is beyond the scope of the study. KO mice also exhibited significant reduction of freezing behavior (Fig. 8 D, p<0.001). To assure there was not anosmia involved in this test, in a separate experiment, we set n=3 introducing double amount of cat litter and observed no differences between the groups that the WT and KO mice spent more than 90% of time freezing. The subjective impression of the experimenters observing the behavior was that the subtle odor stimulus from the container was less salient for the PACAP-deficient mice in that they exhibited enhanced exploratory movements, instead of avoiding the stimulus source and/or freezing, as did the WT mice.

The *fos* expression in KO mice was significantly reduced in olfactory areas, the mitral layer and glomerular layers, as well as higher predator-odor processing centers, for instance, the medial (MeA) and central amygdala (CeA), the lateral septum (LS), the anterior cingulate area (AAC) and prelimbic (PL) and infralimbic (IL) cortical areas, the bed nucleus of stria terminalis oval nucleus (BNSTov); the hindbrain parabrachial complex (PB), which provides the main upstream PACAP signaling source especially for BNSTov and CeA (http://connectivity.brain-map.org/) for the above cognitive centers, and the ventromedial hypothalamic nucleus (VMH), which control the aggressive behavior (Fig. 8 E - G), as well as the cerebellar cortex that we found reduced *fos* expression in the granule cell layer of the aniform lobule, which is reported to influence the limb movement control (Manni and Petrosini 2004, Zhu, Yung et al. 2006). In contrast, KO mice had increased *fos* expression in the motor cortex layer VI and V (Fig. 8 E, H and J).

We assessed *fos* expression in the PACAP/PAC1 system including both glutamatergic and GABAergic neurons in response to odorant exposure, using DISH technique and VGLUT1, VGLUT2 and VGAT mRNAs probes to identify the glutamatergic and GABAergic neurons. Surprisingly, we observed a sharp reduction in the abundance of three vesicular transporters in the main PACAPergic nuclei we described *vide supra* (Fig. 9). This reduction was observed both as reduced abundance of the ISH staining at the single cell level and the density of expressing cells in the given region (Fig. 9 E).

## Discussion

Here, we describe in detail the overall topographical organization of expression of PACAP mRNA and its predominant receptor PAC1 mRNA, and their co-expression with the small molecule transmitters, glutamate and GABA, using VGLUT1-mRNA, VGLUT2 mRNA, and VGAT mRNA, as obligate markers for glutamatergic and GABAergic phenotypes, respectively, in mouse brain. A highly sensitive dual in situ hybridization method (RNAscope 2.5HD duplex detection) was used and corroborated with Allen Brain Atlas ISH data throughout brain and with single-cell transcriptomic data for cortex, with extrapolation to brain regions not yet analyzed in this fashion in the mouse. We report glutamatergic or GABAergic identity of 160 PACAP-mRNA expressing cell groups as indicated by co-expression of corresponding vesicular transporters. All PACAP-mRNA expressing neurons studied and most of their neighboring cells co-express PAC1 mRNA, indicating that the PACAP/PAC1 pathway, beside of using classical neurotra nsmission through axon innervation and transmitter release, likely employs autocrine and paracrine mechanisms for signal transduction.

With the information obtained by PACAP/PAC1 expression and co-localization with small-molecule transmitters, we explored the hypothesis that the broad brain distribution of PACAP may reflect a more specific physiological function at a systems level. We examined the distribution of PACAP/PAC1 expression within specific sensory input-to-motor output pathways passing through the cognitive centers mentioned in these studies, in which most of the hubs analyzed used PACAP>PAC1 signaling within the context of glutamate/GABA neurotransmission. This systematic analysis has revealed several possible PACAP-dependent networks involved in the highest levels of motor control, *i.e*. the hypothalamic pattern initiator and controller system, intermediate (or intervening) levels of somatosensory information processing, and the complex cognitive and behavioral state control for behavior.

### Newly identified PACAP expressing cell groups in mouse brain suggesting circuit-level function

The high-resolution DISH experiment and analysis showed that although the hypothalamus has the highest group-density of PACAP mRNA-expressing subpopulations in the brain, it should not be considered as mainly hypothalamic neuropeptide. Our data revealed a newly identified PACAP mRNA-expressing neuronal populations with mainly VGLUT1 mRNA expression, derived from both *cortical plate* and *brain stem*. Interestingly, most of them co-expressed calcium binding protein calretinin mRNA. The prominent regions/nuclei of this group are: derived from cortical plate: MOB, AOB, AON, TT, PIR, NLOT, COAa from olfactory areas, hippocampus CA3c subset of pyramidal neurons in the ventral pole, dentate gyrus mossy cells, and posterior amygdalar nucleus; bed nucleus of anterior commissure derived from cortical subplate; pontine grey, Koelliker-Fuse of prabrachial complex, nucleus of lateral leminiscus within pons; nucleus of tractus solitarius, dorsal and ventral vestibular nucleus and superior olivary complex lateral part.

### Bed nucleus of anterior commissure (BAC): a prominent yet chemoanatomically ill-defined PACAP-expressing nucleus

We described this nucleus as a newly identified major PACAP-mRNA expressing nucleus (section 3.4, Fig. 5). Regarding the phylogenetic classification of the BAC, some authors have argued that “the septum (within striatum) is divided into the lateral, medial, and posterior septum (LS, MS and PS, respectively); and the PS is further subdivided into the triangular septum (TS) and the bed nucleus of the anterior commissure (BAC) (Risold 2004). However, the Allen Brain Map classifies the BAC as a pallidum structure (https://portal.brain-map.org/). Although literature on BAC connectivity is sparse, LS is identified as one of the input to the BAC (Swanson, Sporns et al. 2016) and the medial habenula is one of the reported target region, with implication in the control of anxiety and fear responses (Yamaguchi, Danjo et al. 2013), the connectivity and function of this glutamatergic-PACAP nucleus is largely elusive. Hence, the chemical identification of this nucleus opens new opportunities for generation of animal models using optogenetic /chemogenetic tools to discern the role of this structure for circuit and behavior.

### A novel transcriptomically distinct pyramidal subpopulation in ventral hippocampal CA3c is well-placed for modulation of the predator threat response

The hippocampus is typically described in the context of the tri-synaptic circuit. The tri-synaptic circuit is composed of three sequential glutamatergic synapses: perforant path axons of layer II neurons in entorhinal cortex project to the outer two-thirds of the dentate gyrus molecular layer, the location of the distal granule cell dendrites; mossy fiber axons of granule cells project to proximal dendrites of area CA3 pyramidal cells; and the Schaffer collateral axons of CA3 pyramidal cells project to stratum radiatum of CA1, where the apical dendrites of area CA1 pyramidal cells are located (Amaral and Witter 1989) (Fig. 4-E). However, this trisynaptic circuit seems insufficient to explain the ventral hippocampus observations about the CA3c pyramidal VGLUT1/PACAP mRNAs co-expressing neurons. Hence, this is a surprising finding since, as discussed by Scharfman (2007), ventral CA3 may be a point of entry that receives information, which needs to be “broadcast,” such as for stress responding, whereas the dentate gyrus may be a point of entry that receives information with more selective needs for hippocampal processing (Scharfman 2007). It has been reported that the CA3c pyramidal cells possess collaterals that project in the opposite direction to the tri-synaptic circuit, “back” to the dentate gyrus, by either direct innervation of the mossy cells and GABAergic interneurons in the hilus, or to the granule cell layer glutamatergic and GABAergic neurons (Scharfman 2007). Those targeted cell types expressed strongly PAC1 mRNA (Fig. 2, A and A’). A hypothetical circuit modified from literature (Scharfman 2007), including this newly identified subset of CA3c PACAP expressing cells, is presented (Fig. 4-F). Hippocampal formation has been suggested to act not as a unitary structure, but with the dorsal (septal pole, DH) and ventral (temporal pole, VH) portions performing different functions (Moser and Moser 1998). Their argument was based on three data sets. First, prior anatomical studies indicated that the input and output connections of the dorsal hippocampus (DH) and ventral hippocampus (VH) are distinct (Swanson and Cowan 1977). Second, spatial memory appears to depend on DH not VH (Moser, Moser et al. 1995). Third, VH, but not DH, lesions alter stress responses and emotional behavior (Henke 1990). Here, we contribute with a fourth element, which is the subset of CA3 pyramidal neurons, located in the temporo-ventral pole of the hippocampal formation, transcriptomically distinct from the rest of CA3 pyramidal neurons, by expressing VGLUT1, PACAP and calretinin mRNAs, which the latter two elements are absent in the septo-dorsal pole.

The VH has attracted attention in odor studies due to dense reciprocal connections to the MeA and to other amygdalar nuclei such as cortical nucleus that receives input directly from main olfactory system (Scalia and Winans 1975) (Fig. 6). The VH also project to AOB and the piriform cortex, a major target of MOB (Shipley and Adamek 1984). It was reported that rats with VH lesions exhibited deficit in freezing and crouching when exposed to cat odor (Pentkowski, Blanchard et al. 2006). Another study in mice exposed to coyote urine showed that VH lesions impaired avoidance and risk assessment behaviors (Wang, Fraize et al. 2013). In the experiments reported here, PACAP-deficient mice (without CA3c of VH expressing PACAP), yielded similar behaviors, suggesting this cell subpopulation may contribute to optimal predator odor processing using PACAP as co-transmitter to coordinate the predator threat response.

### The autocrine/paracrine and neuroendocrine nature of PACAP>PAC1 signaling and glutamate and GABA vesicular transporter expression

The analysis of our data showed an impressive overlapping of PACAP mRNA with PAC1 mRNA at both single cell and regional levels (Fig. 3), suggesting that the PACAP/PAC1 pathway uses *autocrine* and *paracrine* mechanisms in addition to classical neurotransmission through axon innervation and transmitter co-release (Hokfelt, Johansson et al. 1984, Zhang and Eiden 2019). That is, besides sending PACAP-containing axons to innervate regions where PAC1 is strongly expressed, PACAP may be released through soma and dendrites, to bind PAC1 expressed on the same and neighboring cells, to prime the neuron for optimum function (Leng 2018). We still know relatively little about the possible functional interactions and control of release of PACAP>PAC1 signaling in this aspect. However, the observation reported in this study about down-regulation of vesicular transporter mRNAs in PACAP knockout mice, in regions/cell populations, which were normally PACAP mRNA-expressing in wild-type mice, provides one possible example of such a function for PACAP. Activity-dependent regulation of both glutamate and GABA vesicular transporter synthesis and membrane insertion has been reported (De Gois, Schafer et al. 2005, Erickson, De Gois et al. 2006, Doyle, Pyndiah et al. 2010). PACAP signaling most commonly leads to a net increase in neuronal excitability through modulation of intrinsic membrane currents and transiently increasing intracellular calcium concentration (Johnson, May et al. 2019). Decreased vesicular transporter mRNA expression would be expected at the cellular level to decrease transmitter quantal size, thus decreasing excitatory/inhibitory postsynaptic currents (EPSCs and IPSCs respectively) (Billups 2005), and is consistent with our behavior data. PACAP absence in KO mice resulted in a behavioral hypoarousal response to moderate predator-odor stimulus. The rather profound loss of vesicular transporter mRNA upon complete PACAP deficiency may reflect a modulatory role for PACAP, in wild-type animals, on vesicular transporter mRNA (and protein) abundance within a more restricted, but functionally relevant, range depending on the level of PACAP release and autocrine signaling.

PACAP-deficient mice show hyperlocomotion and abnormal gait, as well as bouts of repetitive jumping (data not shown, but see Hashimoto, Shintani et al. 2001, Gaszner, Kormos et al. 2012, Hattori, Takao et al. 2012). In 1930, Hinsey, Ranson and McNattin reported a visionary experimental result, done in cats and rabbits about the rôle of the hypothalamus in locomotion (J. C. Hinsey 1930). Quoting Swanson’s interpretation of this early experiment: “when the central nervous system is transected roughly between the mesencephalon and diencephalon, the animals displayed no spontaneous locomotor behavior. They remain immobile until stimulated. On the other hand, animals with transection roughly between diencephalon and telencephalon (or completely removed the cerebral hemispheres, and the thalamus) display considerable spontaneous behavior. In fact, they can be *hyperactive* when the transection is a bit caudal, but cannot spontaneously eat, mate, or defend themselves. Notably, these last three functions are preserved at a primitive level when the transection is just a slightly more rostral”. This evidence, combined with the selective lesions or stimulations of the hypothalamus, suggests that the ventral half of the diencephalon (hypothalamus) contains neural mechanisms regulating setpoints for locomotor and other classes of motivated behavior” (Fig. 8.9 (Swanson 2012)). As reported here, the major cell groups in the ventral diencephalon, associated with behavioral state (Fig. 7, left column) and behavioral control (Fig. 7, right column) are all using PACAP/PAC1 signaling for co-transmission. Hence, movement pattern/initiation disorders should thus be expected when PACAP is knocked out. Moreover, we found the marked reductions of VGAT mRNA in Purkinje cells (PC) of the ansiform lobule of the cerebellum, as well as significant fos expression reduction in the granule cells (Fig. 9 panels Ds), the main excitatory input of the PC (Manni and Petrosini 2004). The reduced VGAT availability in PC in the lateral lobules of the cerebellum, mainly coordination the movement patterns of the limbs (Manni and Petrosini 2004, Sakayori, Kato et al. 2019), necessarily weakens the inhibitory influence on the excitatory output of the deep cerebellar nuclei which make ascending projections to several forebrain and midbrain structures, including hypothalamus (Zhu, Yung et al. 2006). In Fig 10, we propose a model based on the analysis of this study to provide a rationale for hyperlocomotion and hypoarousal behavior observed in PACAP deficient mice. Under normal conditions, the hypothalamic motor initiator / controller centers (Fig. 7, longitudinal PACAP cell groups in hypothalamus) are modulated, for instance, by direct excitatory (e.g. from deep cerebellar nuclei, DCN), indirect excitatory (through disinhibition of GABAergic projections from BNST, CeA to GABAergic neuronal circuit of hypothalamus) and inhibitory (from lateral septum) influences (Swanson 2012) (Fig. 10-A). We showed in this study that PACAP deficiency caused sharp downregulation of both vesicular glutamate and GABA transporters in the key nuclei (PACAP-expressing in wild type mice), which include cerebeller Purkinje cells exerting inhibitory input to DCN, prefrontal cortex, with reduced VGLUT1 expression (Fig. 9), which would result in a deficient activation of striatum and pallidum cognitive structures, sending GABAergic input to the hypothalamic centers, through direct inhibition or indirect excitation through disinhibition mechanisms, for locomotor patterning and initiation. Unbalanced excitation/inhibition occurring in this hypothalamic region may result in hyperexcitation to motor pathways ending in primary motor cortex (Fig. 8, higher fos expression in primary motor cortex in KO subjects), which command the spinal motor neurons for muscle activities (Fig. 10-B).

## Summary

Voluntary behaviors are mediated by sensory input processed via cognition and brain state to control motor output. The chemoanatomical characteristics of excitatory and inhibitory neurotransmitter co-expression in PACAP- and PAC1-expressing neurons and circuits reveal functionally specialized PACAP-PAC1 neuronal hubs, with specific phylogenetic origins, that indicate their predominance in the sensory, cognitive and brain state to motor control systems. Overall, this data set reveals a connection between PACAP expression in specific brain circuits and the glutamatergic and GABAergic chemotypes of PACAP containing neurons within those circuits, which could serve as a tangible framework for understanding how peptide-small molecule co-expression might function both synaptically and transcriptionally in CNS neurons.

## Materials and methods

### Mice

The following male mouse strain/line were used in this study: C57BL/6N and C57BL/6N PACAP deficient, age 8-10 weeks, generated through an in-house breeding program in accordance with NIH guidelines and standards. Generation of PACAP knock-out mice have been described previously (Hamelink, Tjurmina et al. 2002). All experiments were approved by the NIMH Institutional Animal Care and Use Committee (ACUC) and conducted in accordance with the NIH guidelines.

### In situ hybridization (ISH): RNAscope single and duplex (DISH) procedure

Mice were deeply anesthetized with isoflurane, decapitated and whole brains were removed, embedded in OCT medium and rapidly frozen on dry ice. Serial brain sections (12 μm thick) were cut on a cryostat (Leica CM 1520), adhered to SuperFrost Plus slides (ThermoScientific). RNA probes were designed and provided by Advanced Cell Diagnostics (Hayward, CA): Mn-Adcyap1 (gene encoding the protein PACAP), Mn-Adcyapr1r (gene encoding the PACAP receptor 1), Mn-Slc17a7 (gene encoding the protein VGLUT1), Mn-Slc17a6 (gene encoding the protein VGLUT2), MnSlc17a8 (gene encoding the protein VGLUT3), Mn-Slc32a1 (gene encoding the protein VIAAT), Mn-Vipr1 (gene encoding the VPAC1 receptor), Mn-Vipr2 (gene encoding the VPAC2 receptor), Mn-Sst (gene encoding somatostatin), Mn-Pvalb (gene encoding palvalbumin), Mn-Crh (gene encoding corticotropin releasing hormone), and Mn-Fos (gene encoding the protein cFos). All staining steps were performed following the protocols provided by the manufacturer for chromogenic detection of mRNA in fresh frozen tissue samples. Stained slides were examined with both light microscope with digital camera and were image-captured with 20X objective with ZEISS Axio Scan (Carl Zeiss Microscopy, Thornwood, NY from Systems Neuroscience Imaging Resource, NIMH-IRP, NIH) and images for sections from each animal were organized and converted to TIF files with BrainMaker (MBF Bioscience, Williston, VT).

### Chemo-anatomical analysis and circuit identification

Anatomical nomenclature and regional delineation were done according to Allen Mouse Brain Atlas (www.allenbrainatlas.org) corroborated with Paxinos and Franklin’s the Mouse Brain in Stereotaxic Coordinates. The PACAP/PAC1 mRNAs distribution mapping and semi-quantification and their co-expression with glutamate/GABA transporters mRNAs were corroborated with the Allen brain atlas https://mouse.brain-map.org/. For schematic circuit chartings of figures 6, 7 and 10, Swanson’s rodent flat maps downloaded from http://larrywswanson.com/?page_id=164. was used. Sensorimotor circuit hubs and connectivity of figures 6 and 10 are based on literature cited in the results section as well as consulting the website https://sites.google.com/view/the-neurome-project/connections/cerebral-nuclei?authuser=0.

### Behavior experiment

**Subjects:** C57BL/6N wild type and PACAP deficient male mice, age 8-10 weeks, n=9 (N=18) were used. They were group-housed (4/cage) in a room kept on a controlled light-dark cycle (light on 7:00 am and off 7:00pm) under constant humidity and temperature conditions.

**Odor stimuli:** cat urine material was collected from domestic cat litter, where multiple male and female cats used the same box to urinate.

**Modified open-field box for odor exposure and fos expression assessment:** a custom-made wooden box (28cm x 28cm x 28 cm, with a sliding glass lid) was used (see figure 8-A). The box was positioned inside a low-noise suction hood, then in a lidded container, cat urine containing litter material was introduced into the box (figure 8-A). After introducing the subject, the lid of container was opened through a string and the box was closed to avoid the odor to escape. Video recording of the animal behavior was made during a 10 min period, after which animals were returned to its home cage for 30 min and then euthanized by cervical dislocation and brains rapidly removed and processed for dual in situ hybridization (DISH) using the RNAscope 2.5 HD Duplex Assay. The expression of *fos* mRNA within PACAP, PAC1, VGluT2 and VGAT-positive cells.

**Behavior assessment:** mouse behavior during the 10 min of cat urine exposure was recorded by an overhead camera. Behavioral scoring was performed off-line with *blind* analysis (performed by ECG, medical student in laboratory rotation, see acknowledgement. Freezing behavior was assessed in the first 5 min lapse, giving a score every 5 sec when the subject exhibit immobility with piloerection. Complete cycle of sniff>retreat was defined as the movement of approach the containing and immediately retreat beyond the quadrant I (QI, figure 8) while incomplete sniff > retreat cycles were defined when the subject approached the container and stayed in the proximity or retreat only a short distance within the same quadrant (Q1). Mice spatial displacement maps were produced on-screen by the analyst with aid of the software PowerPoint > shape format > curve tool (Microsoft Office). The experimental arena was represented in a slide with a square containing 16 small squares. The flask containing the cat litter is symbolized with a filled circle. The computer aided manual drawing was visually guided by the video recording under the strict criterion that the analyst makes a click in the corresponding position of the map for each *end of a lineal movement* of the mouse. The jumping behaviour is only symbolized with “zig-zags”. Four of mice exhibited this behavior and due to the aim of this study - the place preference assessment - these subjects were discarded for the Heat map. PowerPoint traces where skeletonized with ImageJ and movement traces from n=5 animals were used. The 2D movement heat-map was produced with MATLAB R2016b, for each time the mouse passed through a point in space, the value of 1 was assigned. Using the MATLAB “color dispersion function” (PSF) and “color map” functions a heat map was constructed. The trajectories of the 5 mice in each group were superimposed, then using the ‥bwarea‥ function, a count of the pixel quantity per quadrant was performed, being Q1 the container located quadrant (clockwise numbering). Thereby, each pixel in the image had a different value and color according to the number of times the mouse was at that point. The number of pixels per quadrant for mice in the KO group was compared with mice in the WT group following one-way ANOVA.

### Data analysis

GraphPad Prism 7.0 was used to perform Student t-tests and one-way ANOVA for evaluation of statistical differences between groups, levels are indicated as follows: *P < 0.05, **P < 0.01, ***P < 0.001 Unless otherwise indicated, values are reported as mean ± SEM.

## Competing interest

The authors declare that no competing interest exist.

## Funding

see page 32

## Acknowledgements

We apologize to colleagues whose contributions may have been overlooked in citation of the relevant literature for this report. We thank Manuel Hernández for producing the modified open-field box for the predator odor test, to students Enrique C. Guerra, Anil K. Verma and Sean Sweat for technical assistance and to Angel Fermín Barrio-Zhang for the drawing of the experimental design. LZ was a Fulbright visiting scholar to NIMH-IRP, NIH. LZ and RAB were on sabbatical stay hosted by LEE (NIMH), supported by PASPA-DGAPA-UNAM fellowships. LZ also would like to thank Peter Somogyi and Rafael Lujan for hosting her academic research visits and discussions when part of this manuscript was developed. We thank Peter Somogyi and Rafael Lujan for critical reading and comments on an early version of this manuscript.

## Grants

UNAM-DGAPA-PAPIIT-IN216918, IG200121 (LZ) & CONACYT-CB-238744 (LZ), CB-283279 (RB) and NIMH-IRP-1ZIAMH002386 (LEE).

## Supplementary information (three figures and one table)

## Notes

### Competing Interest Statement

The authors have declared no competing interest.

### Summary of Updates

1) Figures 7 and 10: diagrams were edited to correct some minor colored shapes errors. 2) Figure 10's legend was edited due to missing abbreviations definitions. 3) Some spelling errors in the main text were corrected.

