## Supplementary information for "Behavioral role of PACAP signaling reflects its selective distribution in glutamatergic and GABAergic neuronal subpopulations" (from another version of this preprint: version 1, 2020-08-01)

*Running title:*

*PACAP/PAC1 cell-types and circuit interaction*

**Supplementary figure 1 (1/7):** Comprehensive DISH mapping of PACAP co-expression with VGLUT1, VGLUT2 and VGAT throughout mouse brain reveals an extensive distribution and diversity of cell types.

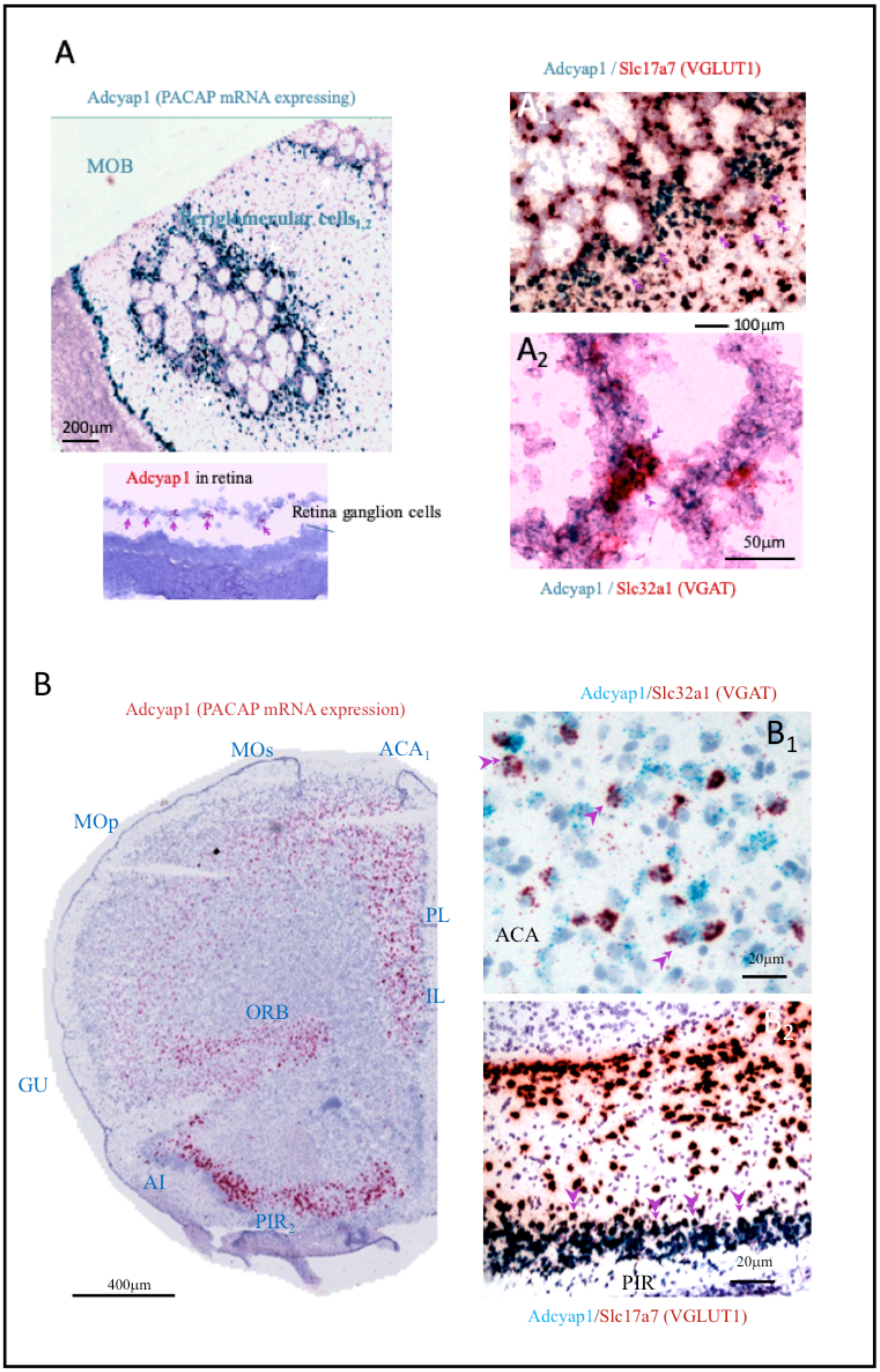

**Supplementary figure 1 (2/7)**

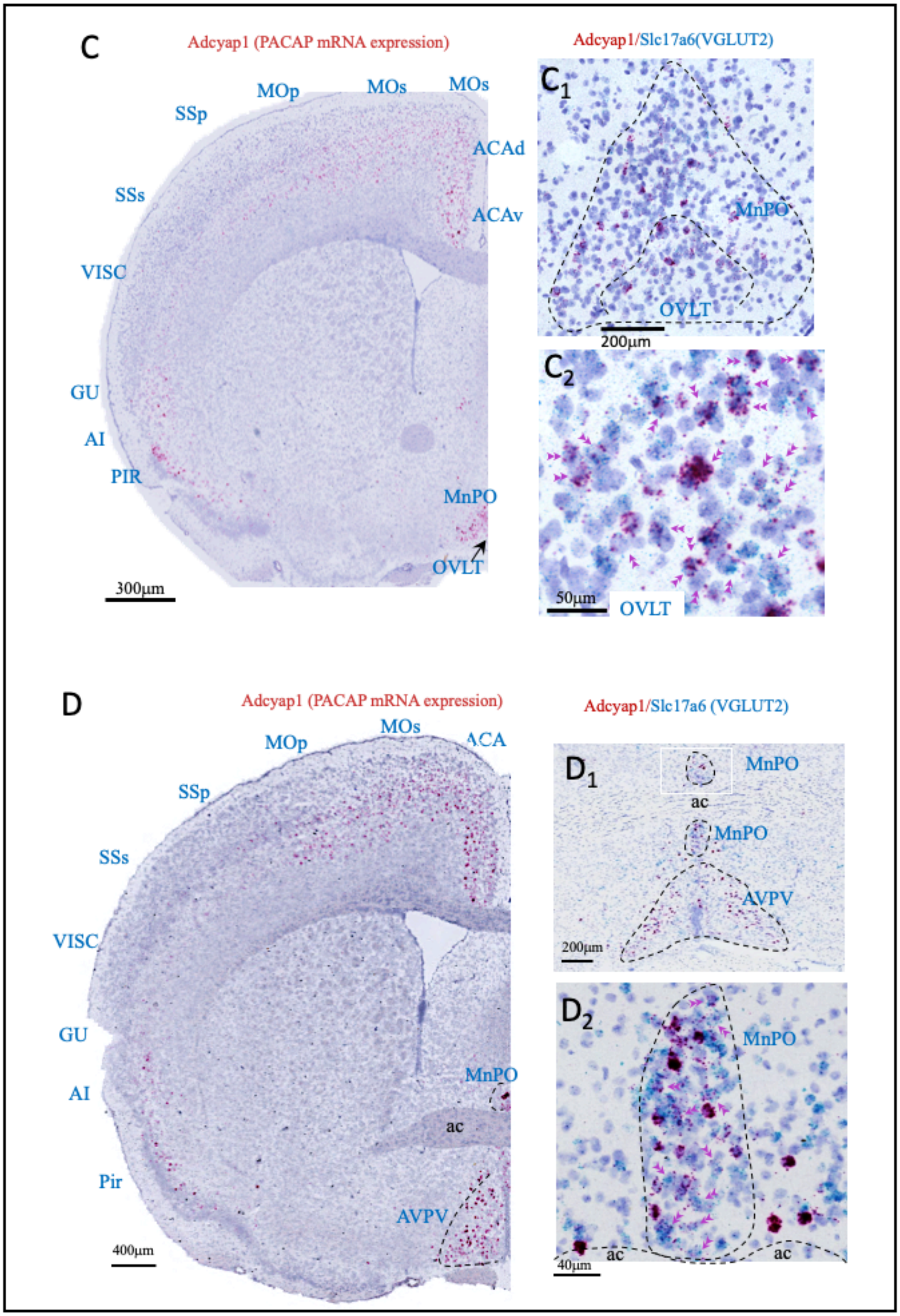

**Supplementary figure 1 (3/7):**

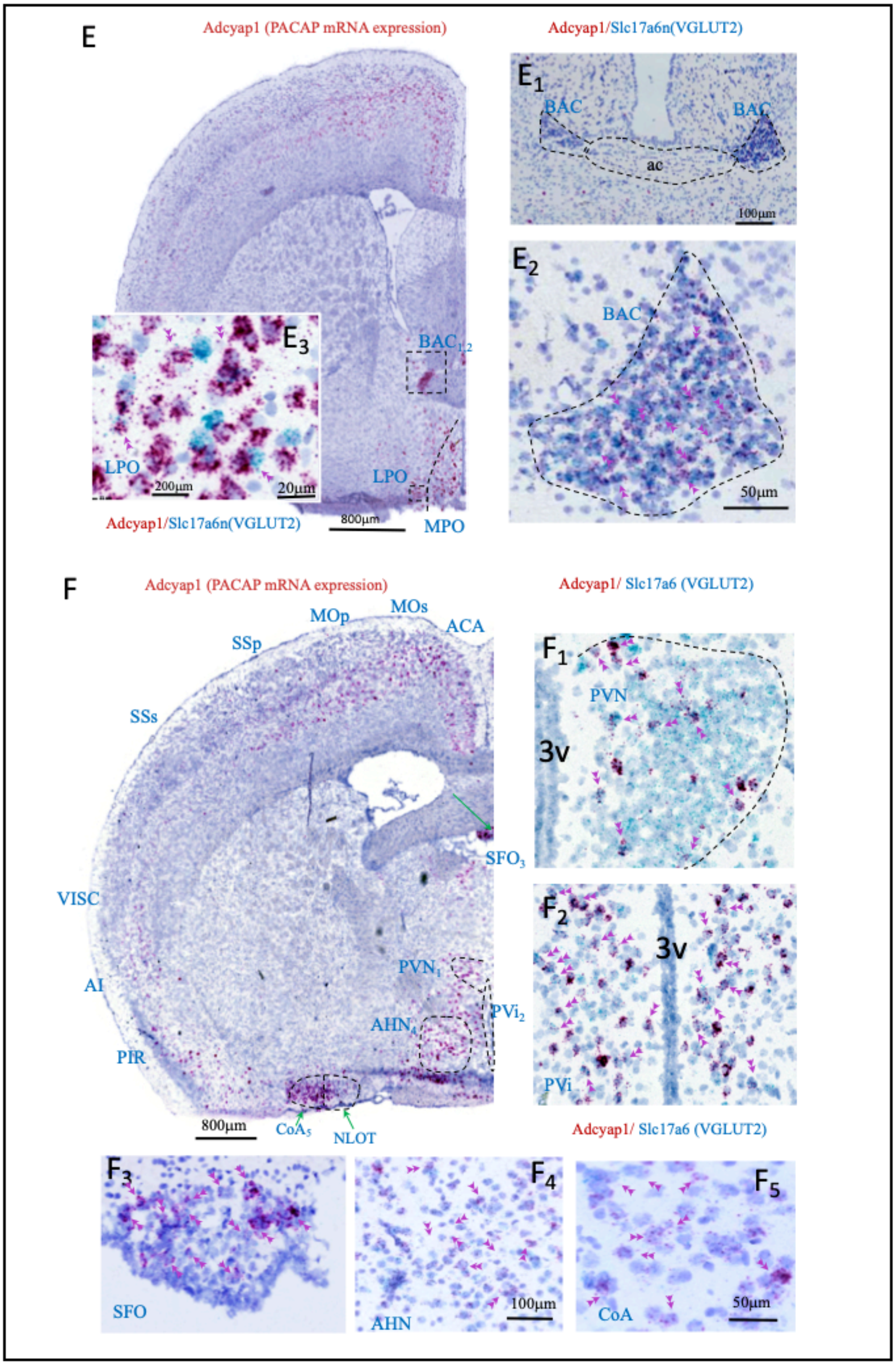

**Supplementary figure 1 (4/7):**

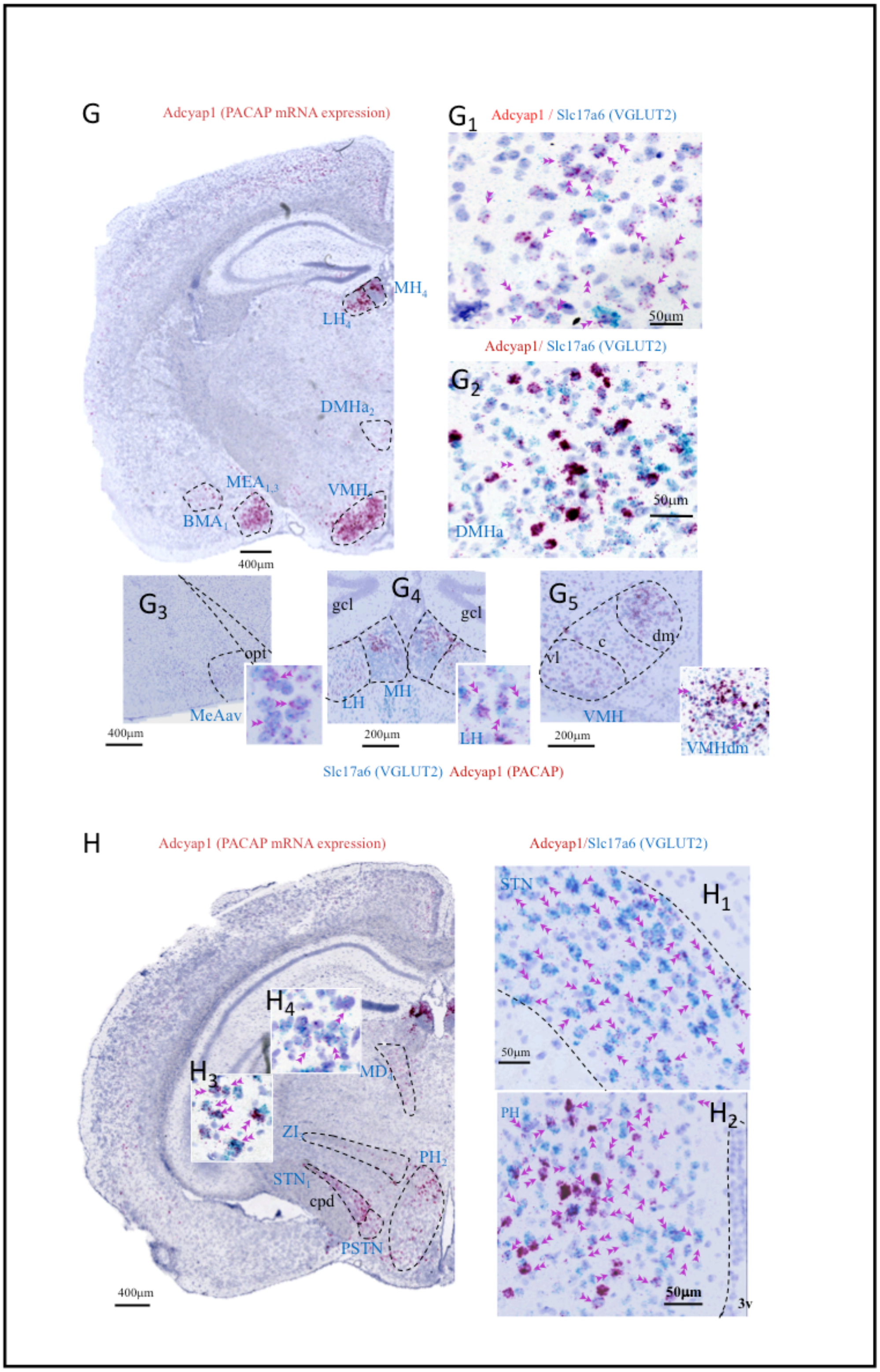


**Supplementary figure 1 (5/7):**

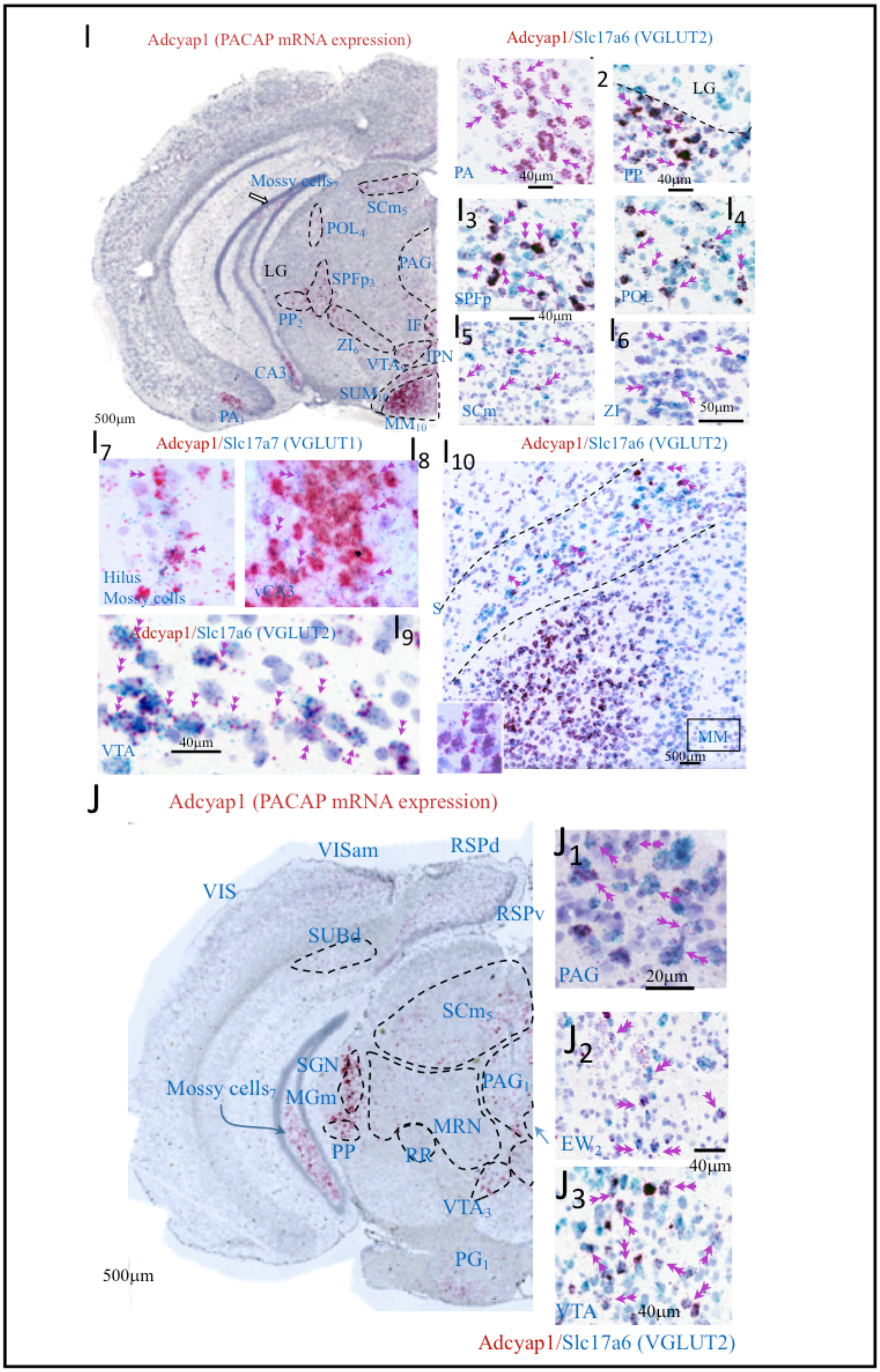
I
Adcyap1 (PACAP mRNA expression)
Adcyap1/Slc17a6 (VGLUT2)
Mossy cells7
SCm5
POL4
LG
SPFp3
PAG
PP2
IF
ZI6
VTA
IPN
CA38
SUM
PA1
MM10
500µm
PA
40µm
PP
I3
I4
SPFp
POL
I5
I6
SCm
ZI
50µm
I7
Adcyap1/Slc17a7 (VGLUT1)
I8
I10
Hilus
Mossy cells
vCA3
Adcyap1/Slc17a6 (VGLUT2)
I9
VTA
MM
J
Adcyap1 (PACAP mRNA expression)
VISam
RSPd
VIS
RSPv
SUBd
SCm5
SGN
Mossy cells7
MGm
PAG1
MRN
PP
RR
VTA3
PG1
500µm
J1
PAG
20µm
J2
EW2
J3
VTA
Adcyap1/Slc17a6 (VGLUT2)

**Supplementary figure 1 (6/7):**

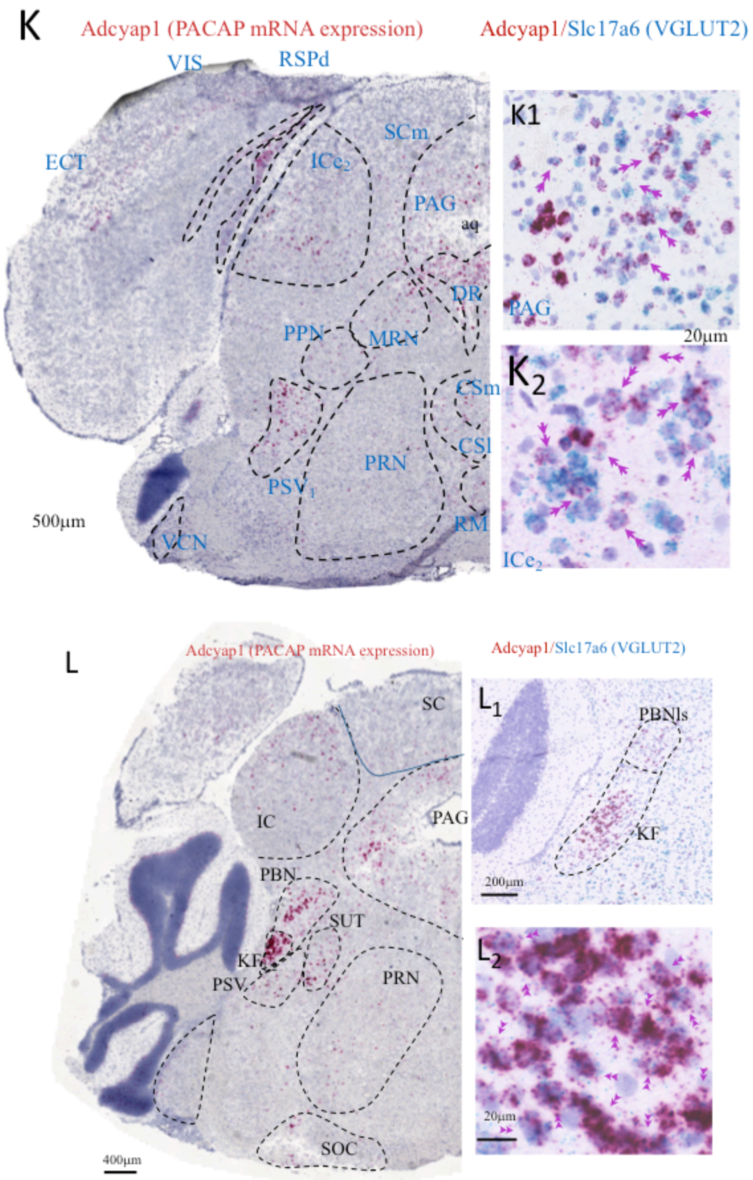

**Supplementary figure 1 (7/7):**

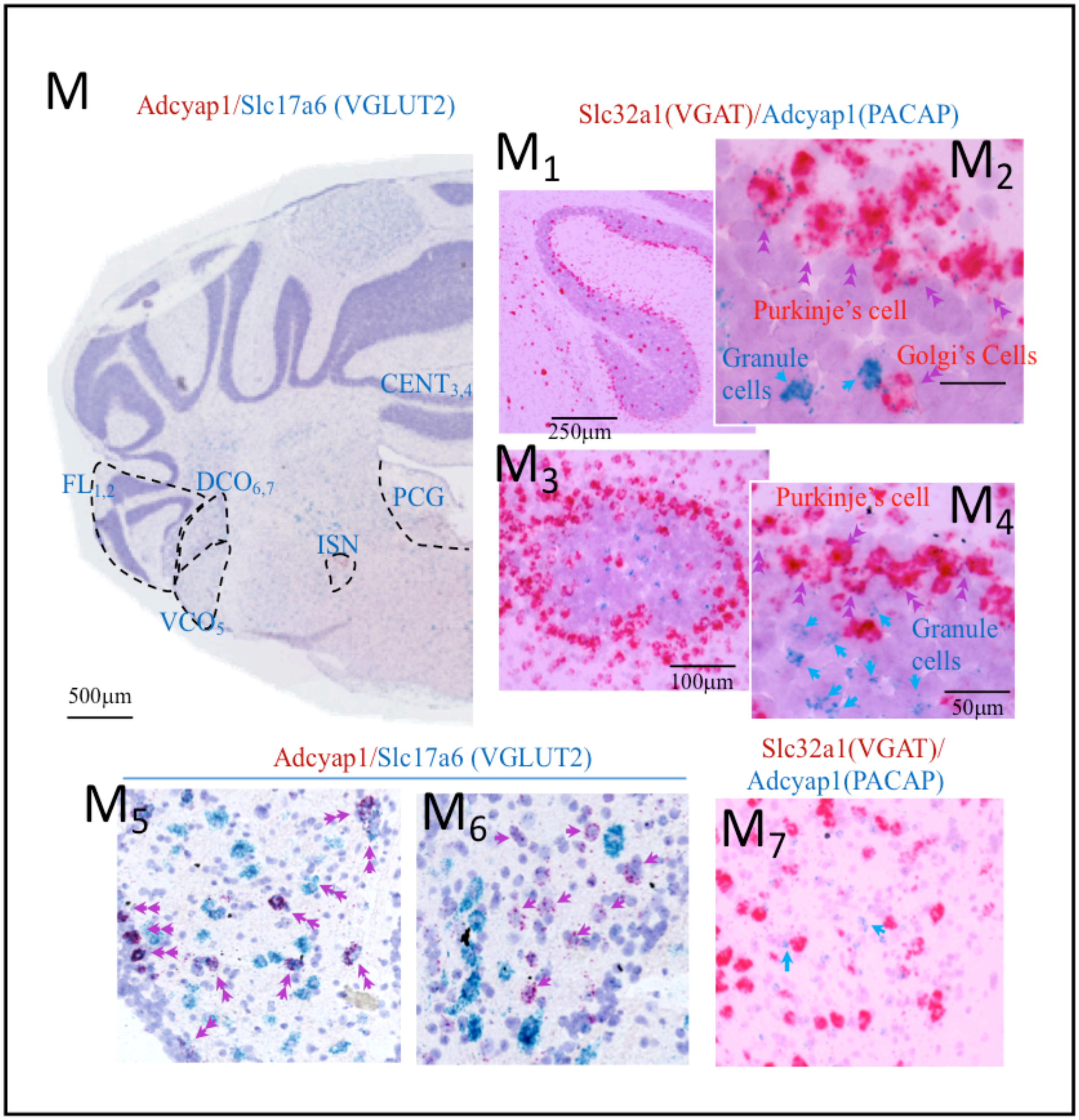


**SI-Fig. 1. Comprehensive DISH mapping of PACAP co-expression with VGLUT1, VGLUT2 and VGAT throughout mouse brain reveals an extensive distribution and diversity of cell types.** Adcyap1, PACAP-mRNA mapping within subpopulations of glutamatergic and GABAergic subpopulations of mouse brain. Scl17a7, Slc17a6, Slc17a8 and Slc32a1: mRNAs encoding the VGLUT1, VGLUT2, VGLU and VGAT, respectively.

**Supplementary figure 2:**

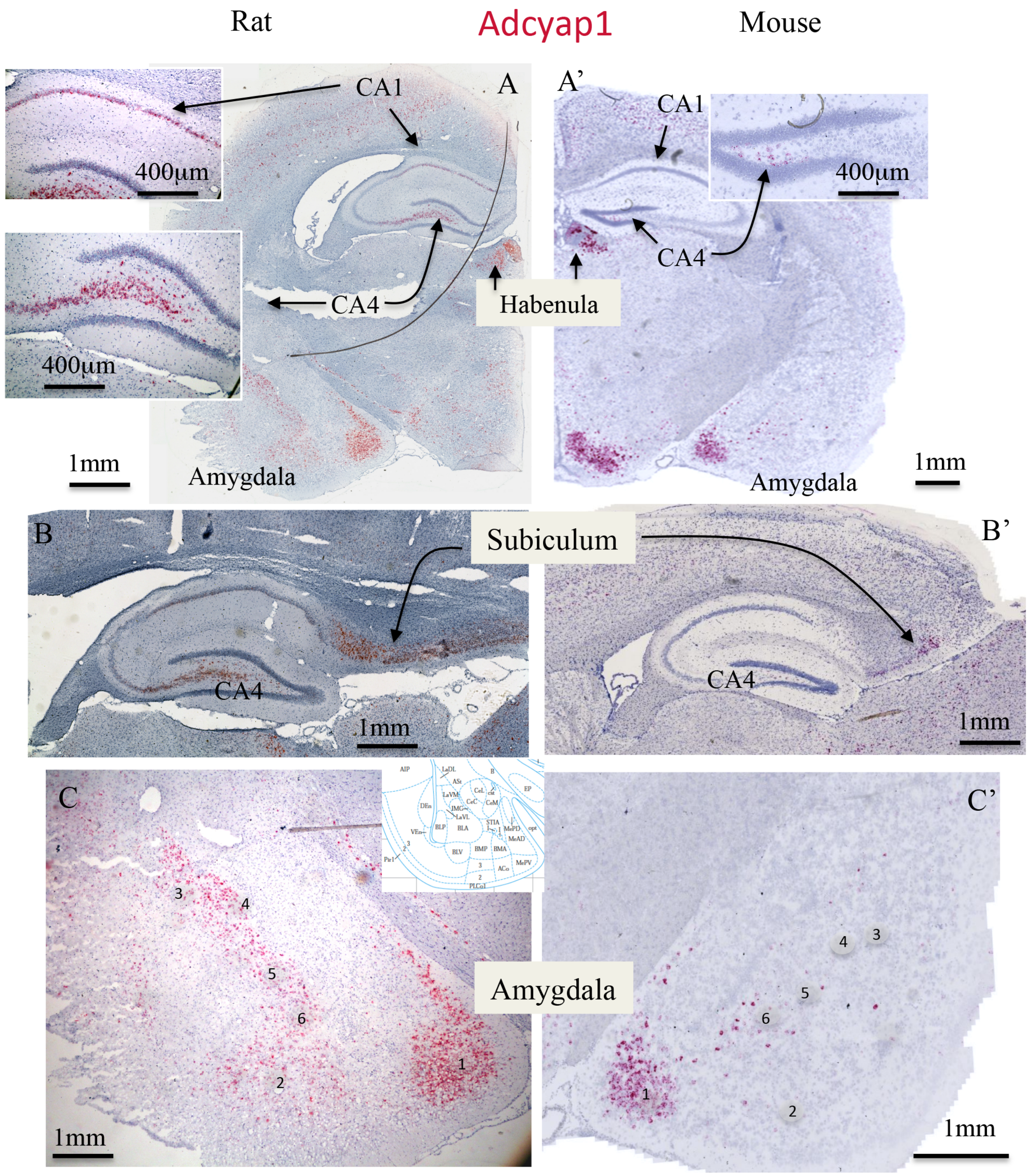


1. corticomedial amygdala CoM; 2. basal amgdala; 3. central amygdala, capsular; 4. central amygdala, lateral; 5. intercalated cells complex; 6. bed nucelus of stria terminalis, intraamygdalar division; 7. dorsal endopiriform nucleus

**Supplementary figure 2: Neurochemical and anatomical divergence between the distribution of PACAPergic circuits in phylogenetically old and new brain areas were compared, revealing possible evolutionary divergence of PACAP function in mouse and rat. PACAPergic cell distribution** within the subcortical regions (such as habenula and hypothalamus are relatively well conserved between the two rodent species. However, phylogenetically new regions we observed more abundant Adcyp1 expression. Here are two limbic regions examined with RNAscope in situ hybridization using Adcyap1 probe (see the method in the next section) as examples. A and A': coronal sections from young adult male rat and mouse showing the conserved patterns of expression in subcortical regions of hypothalamus and habenula, but enhanced expression in rat in hippocampal formation and amygdala complex. Top inset of A shows hippocampus CA1 has high expression of Adcyap1 while in mouse this feature is not seen. Lower inset of A shows abundant expression of CA4/hilus region in rat while few of Adcyap1 expressed cells were observed in dorsal hilus region in mouse. B and B', sagittal sections from rat and mouse showing the increased expression of Adcypa1 in rat subiculum. C and C', comparison in amygdala complex while in rat, 6 regions with abundant expression of Adcypa1 were observed: 1. corticomedial amygdala CoM; 2. basal amgdala; 3. central amygdala, capsular; 4. central amygdala, lateral; 5. intercalated cells complex; 6. bed nucelus of stria terminalis, intra-amygdalar division; 7. dorsal endopiriform nucleus. In the same regiones, in mouse, some scattered Adcyap1 expressed cells can also be seen.

**Supplementary Table.** 1 Density distribution of **PAC1 expressing cells** in cortical regions

| Cortex | | | |
|---|---|---|---|
| **Cg1/Cg2** | PAC1 | **RSG/RSA** | PAC1 |
| **I** | ++++ | I | ++++ |
| **II/III** | +++++ | II/III | +++++ |
| **V** | +++++ | V | ++++ |
| **VI** | +++++ | VI | +++ |
| **M** | | **M&V & S1** | |
| **I** | +++ | I | +++ |
| **II/III** | +++++ | II/III | +++++ |
| **IV** | ++++ | IV | ++++ |
| **V** | ++++ | V | ++++ |
| **VI** | +++ | VI | +++ |
| **S1/S2** | | **AuV/Ect** | |
| **I** | +++ | I | +++ |
| **II/III** | +++++ | II/III | +++++ |
| **IV** | ++++ | IV | ++++ |
| **V** | ++++ | V | ++++ |
| **VI** | ++++ | VI | ++++ |
| **GI/DI** | | **ENT** | |
| **I** | +++ | I | +++ |
| **II/III** | +++++ | II/ | +++++ |
| **IV** | ++++ | III/IV | +++ |
| **V** | ++++ | V/IV | +++++ |
| **VI** | +++++ | | |

Semiquantitative annotations are used here (the percentage of PAC1 expressing cell/total Nissl stained nuclei: “+++”, moderate (60%-40%); “++++”, intense (80%-60%); “+++++”, very intense (>80%).

**Supplementary figure 3**

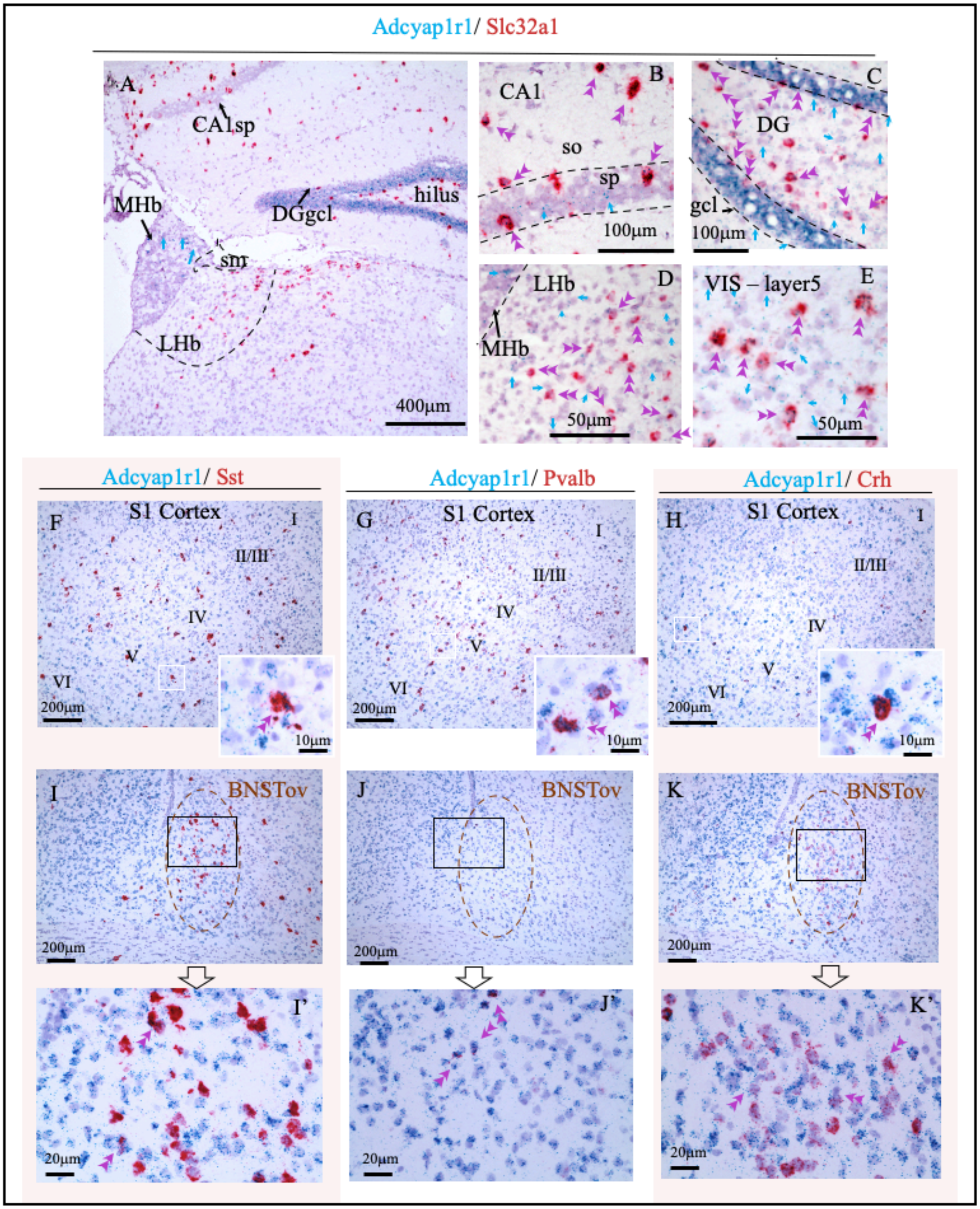

**Supplementary figure 3: Examples illustrating PAC1 mRNA (Adcyap1r1) was highly co-expressed in cortical and subcortical GABAergic neurons.** A: low magnification showing epithalamic habenula and hippocampus with Slc32a1 (VGAT mRNA) expressing cells (red) sparsely distributed. B and C: in hippocampus CA1 (B), both stratum oriens (so) and stratum pyramidale (sp), and dentate gyrus (C), hilus of dentate gyrus (DG) and granule cell layer (gcl), PAC1 mRNA (Adcyap1r1, light blue) was co-expressed with all VGAT mRNA (Slc32a1, red) expressing neurons (double arrowheads). Some principal cells in sp also expressed PAC1 (single blue arrowheads) while in glc, all principal cells co-expressed Adcyap1r1. D. in lateral habenula, all the Slc32a1 cells co-expressed PAC1 mRNA (Adcyap1r1) as well as some no-VGAT expressing cells. E. in primary visual cortex, layer 5, both Slc32a1 expressing and no-expressing cells express Adcyap1r1. F, G and H show the co-expression of Adcyap1r1 (PAC1) with main types of cortical GABAergic interneurons containing somatostatin, parvalbumin and corticotropin releasing hormone (Sst, Pvalb and Crh respectively), in the primary somatosensory cortex (S1). I-VI indicate cortical layers. I, J and K, the same mRNAs co-expression (of F, G, H) in the bed nucleus of stria terminalis, the oval subnucleus (BNSTov). I', J' and K' are amplifications of the corresponding black-squared fields. Double arrow heads indicate some co-expressed cells.
